# Structural Insights into the Integration of Temperature and pH by Sperm Calcium Channel CatSper

**DOI:** 10.64898/2026.04.10.717635

**Authors:** Billy Zhao, Shweta Bhagwat, Juan Ferreira, Dilip K. Swain, Celia Santi, Ziao Fu, Polina V. Lishko

**Affiliations:** Department of Cell Biology and Physiology, WashU Medicine, Washington University in St. Louis, School of Medicine, St. Louis, MO, USA; Center for the Investigation of Membrane Excitability Diseases (CIMED), WashU Medicine, St. Louis, MO, USA; Department of Obstetrics&Gynecology, WashU Medicine, Washington University in St. Louis, School of Medicine, St. Louis, MO, USA; BJC Investigator Program, WashU Medicine, St. Louis, MO, USA

**Keywords:** cationic channel of sperm, CatSper, CatSper1, histidine-rich protein, pH-sensitivity, temperature-gating, evolution, sperm, male fertility, infertility, temperature and voltage-dependence

## Abstract

The cation channel of sperm (CatSper) is a sperm-specific calcium channel essential for male fertility across metazoans. Its activation is tightly restricted to defined physiological contexts, including intracellular alkalinization, membrane depolarization, and elevated temperature. The structural and evolutionary mechanisms underlying this polymodal integration remain poorly understood, in part due to the architectural complexity of CatSper, a ∼15-subunit assembly organized in zigzag arrays along the sperm flagellum. Here we combine comparative genomics across 47 species with AlphaFold3-based modeling and evolutionary sequence–structure analyses to uncover a mechanism for temperature and pH integration. We identify the pore-forming subunit CatSper1 as an evolutionary hotspot exhibiting exceptional divergence in its N-terminal domain. Phylogenetic analysis shows that N-terminal length and histidine enrichment scale with species-specific fertilization temperatures, suggesting adaptive tuning of physicochemical sensitivity. Structural modeling indicates that conserved surface-exposed histidine clusters form inter-complex coupling interfaces between adjacent CatSper assemblies positioned near the dominant voltage-sensing module. Functional validation using electrophysiology and calcium imaging in mouse sperm shows that capacitation-associated partial removal of the CatSper1 N-terminus selectively impairs temperature-dependent activation.

Together, these findings support a model in which temperature-dependent histidine deprotonation modulates supramolecular CatSper assembly to coordinate channel activation.

## INTRODUCTION

The cation channel of sperm (CatSper) is a sperm-specific, calcium ion channel that is essential for male fertility across the evolutionary tree ^1–15^. Originally identified in mice ^1^, CatSper channels are indispensable for hyperactivation-an asymmetric flagellar bending pattern required for successful fertilization-which is triggered by calcium influx through CatSper ^1,8,16^. Because calcium entry must be tightly controlled, CatSper activity is regulated by multiple modalities that ensure it functions only under precise physiological conditions. CatSper is gated by warm temperatures ^17^, membrane depolarization ^6,12,18,19^, and intracellular alkalinization ^6,12,19^, while cytoplasmic acidification potently inhibits its function ^6^.

This multimodal regulation is supported by the structural complexity of CatSper, which forms a large multiprotein complex composed of at least 15 distinct subunits, each encoded by a separate gene ^1,4,7,13,17,20–32^, and is called CatSpermasome ^31^. Each CatSpermasome is further assembled into highly ordered zigzag arrays that form four longitudinal rows running along the entire length of the sperm principal piece ^21^.

Comparative genomic studies have revealed that while CatSper genes are broadly conserved across vertebrates, they exhibit striking lineage-specific patterns of gain and loss, highlighting the channel’s evolutionary diversity ^10,15,20,33^. In mammals such as mice and humans, CatSper is indispensable for male fertility; however, in certain lineages—including birds—the channel has been lost, suggesting the evolution of alternative mechanisms for sperm activation ^15,34^.

The pore of the CatSper channel is formed by the relatively homologous subunits CatSper1, CatSper2, CatSper3, and CatSper4 ^7^. All four are six-transmembrane domain polypeptides encoded by separate genes, and they share a conserved selectivity filter. While most other CatSper subunits are conserved, one in particular-CatSper1-shows exceptional divergence across species, particularly in the length of its N-terminal domain highly enriched in histidine residues.

To explain the multimodal regulation of CatSper, such as its ability to integrate pH and thermal signals to fine-tune calcium entry during sperm hyperactivation, we combined comparative genomics with AlphaFold3 ^35^ prediction and built a model revealing the potential mode of operation of this unique ion channel. According to this model, surface histidine residues in the N-terminal domain of CatSper1 engage in pH-dependent interactions, stabilizing a previously considered disordered domain, and coupling the entire array of CatSper channels. Elevated temperature causes histidine deprotonation, enhancing the coupling and leading to synchronized channel opening. The importance of CatSper1 N-terminal domain for temperature sensitivity was further supported by electrophysiological and calcium imaging recording from murine sperm that underwent partial degradation of N-terminal domain^14,17,36^.

This phylogenetic diversity underscores the importance of CatSper channels not only as key regulators of mammalian reproduction but also as a model for understanding how evolutionary pressures shape reproductive physiology and gamete fitness.

## RESULTS

### Comparative genomics analysis revealed a correlation between N-terminal composition of CatSper1 and optimal fertilization temperature

The core of the CatSpermasome complex is formed by four pore-forming subunits: CatSper1, CatSper2, CatSper3, and CatSper4^7^, while four auxiliary subunits-CatSper β^27^, δ ^13^, ε ^23,37^ and γ ^25^ - extend large extracellular domains that form a canopy over the channel pore^21,31^. All eight CatSper subunits are transmembrane proteins and are largely conserved across species, with the notable exception of CatSper1, which exhibits substantial variation in length (Fig. 1a-b and Fig S1a) and shows a positive correlation between its overall length and species-specific fertilization temperatures (Fig. 1c and S1b). Strikingly, the feature that sets CatSper1 apart is its unusually long N-terminal domain (NTD), whose length also scales with fertilization temperature and is highly enriched in histidine (His, H) residues (Fig. 1d). Interestingly, marine invertebrates such as corals and sea urchins that spawn their sperm directly into sea water, feature truncated NTD barely reaching 120 amino acids (Fig. 1a-d and Table 1). This trend was shared with reptiles that presented a slightly elongated NTD around 127 amino acids. As the core body temperature increased, particularly for internal fertilizers, the length of the domain rose exponentially, reaching ∼500 residues in species such as *Lontra Canadensis* (River Otter) and *Oryctolagus Cuniculus* (European Rabbit).

**Figure 1.**
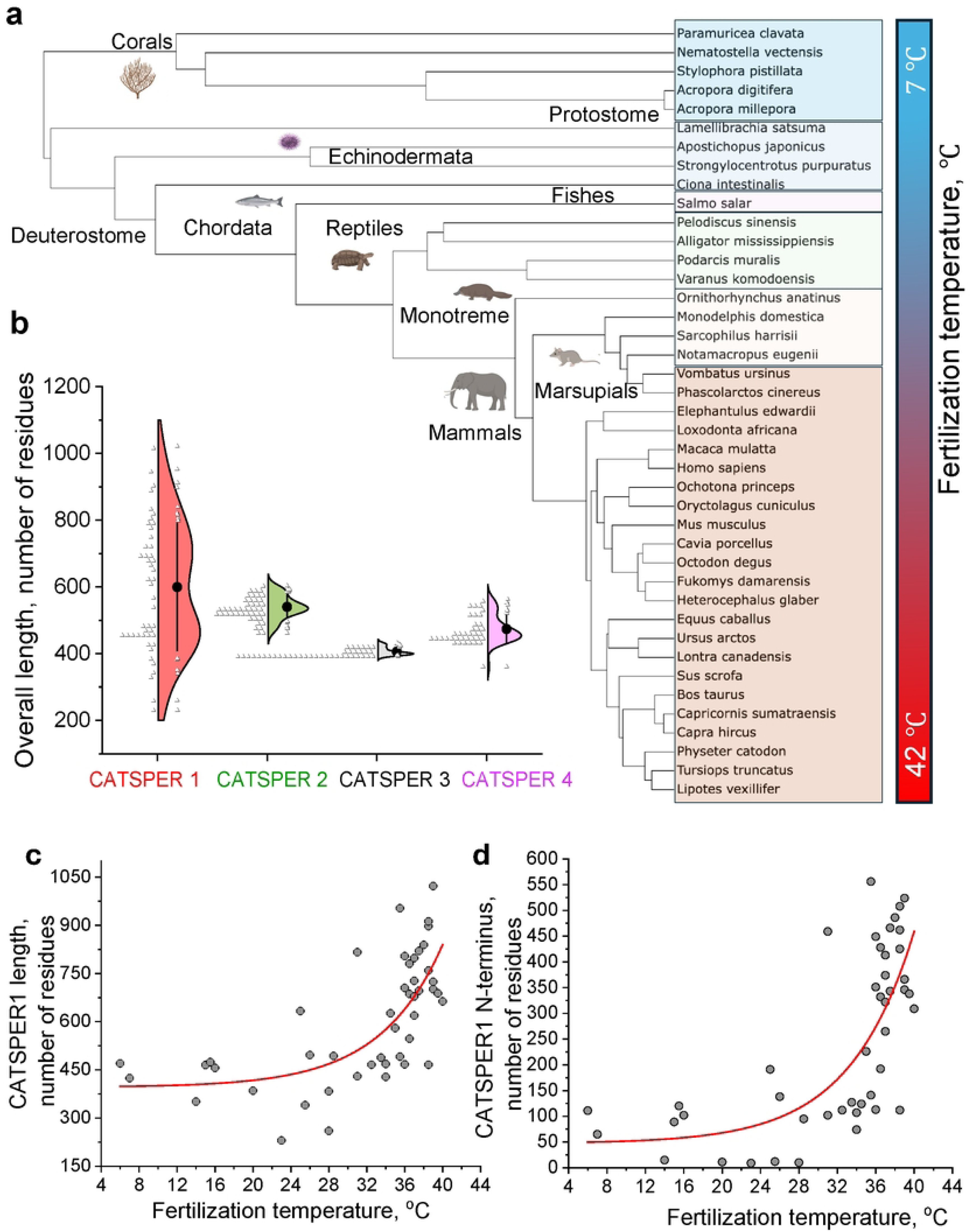
Comparative genomic analyses reveal temperature-dependent divergence of the CatSper1 N-terminus across CatSper-bearing species. (a) Phylogenetic tree of representative species that retain CatSper channels. (b) Comparison of the total amino acid length of the four CatSper pore-forming subunits across the species shown in (a), highlighting pronounced divergence of CatSper1 relative to CatSper2–4. (c) Total CatSper1 length plotted against fertilization temperature for the species shown in (a), revealing an exponential relationship. (d) Length of the CatSper1 NTD plotted as a function of fertilization temperature for the same species, demonstrating a strong positive correlation.

**Table 1.** Species included in this study and their CatSper-related parameters. The lengths of the CatSper1 subunit and its N-terminal domain (NT) are reported as numbers of amino acid residues (aa). Additional columns include the theoretical isoelectric point (pI) of the CatSper1 N-terminus, fertilization temperature in °C, percentage (%) and absolute number of histidine residues (#) within the N-terminus, and the corresponding references for fertilization temperatures for each species. Colored area depict cold-fertilizing species (6–23 °C; blue), intermediate temperatures (25–34.5 °C; green), and warm-fertilizing species (35–40 °C; red).

|  | Species, name | Common name | T, °C | NT, aa | Histidines in NT |  | NT, pI | CatSper1 total length, aa | Citation |
| --- | --- | --- | --- | --- | --- | --- | --- | --- | --- |
|  |  |  |  |  | % | # |  |  |  |
| 1 | *Oncorhynchus mykiss* | Rainbow trout | 7 | 65 | 4.6 | 3 | 10.7 | 424 | 46 |
| 2 | *Salmo salar* | Atlantic salmon | 6 | 111 | 3.6 | 4 | 10.1 | 470 | 47 |
| 3 | *Lamelibrachia satsuma* | Satsuma tube worm | 14 | 15 | 0 | 0 | 4.4 | 351 | 48 |
| 4 | *Ciona intestinalis* | Sea squirt | 15 | 89 | 4.5 | 4 | 9.8 | 465 | 49 |
| 5 | *Strongylocentrotus purpuratus* | Purple sea urchin | 15.5 | 120 | 5.8 | 7 | 6.7 | 474 | 50 |
| 6 | *Apostichopus japonicus* | Japanese sea cucumber | 16 | 102 | 2.9 | 3 | 10.1 | 456 | 51 |
| 7 | *Nematostella vectensis* | Starlet sea anemone | 20 | 11 | 9.1 | 1 | 11.2 | 385 | 52 |
| 8 | *Paramuricea clavata* | Red gorgonian sea fan | 23 | 9 | 33.3 | 3 | 11.0 | 230 | 53 |
| 9 | *Orchesella cincta* | Springtail | 25 | 191 | 2.6 | 5 | 5.3 | 633 | 54 |
| 10 | *Stylophora pistillata* | Smooth cauliflower coral | 25.5 | 12 | 16.7 | 2 | 9.7 | 340 | 55 |
| 11 | *Lingula anatina* | Lingula brachiopod | 26 | 138 | 3.6 | 5 | 6.5 | 496 | 56 |
| 12 | *Acropora digitifera* | Finger coral | 28 | 10 | 10 | 1 | 11.0 | 260 | 57 |
| 13 | *Acropora millepora* | Staghorn coral | 28 | 10 | 10 | 1 | 11.0 | 383 | 58 |
| 14 | *Pelodiscus sinensis* | Chinese softshell turtle | 28.5 | 95 | 1.1 | 1 | 10.0 | 493 | 59 |
| 15 | *Alligator mississippiensis* | American alligator | 31 | 102 | 2.0 | 2 | 11.2 | 430 | 60 |
| 16 | *Heterocephalus glaber* | Naked mole-rat | 31 | 459 | 26.4 | 121 | 6.8 | 816 | 61 |
| 17 | *Ornithorhynchus anatinus* | Platypus | 32.5 | 112 | 3.6 | 4 | 10.7 | 466 | 62 |
| 18 | *Podarcis muralis* | Common wall lizard | 33.5 | 127 | 3.2 | 4 | 11.9 | 488 | 63 |
| 19 | *Vombatus ursinus* | Common wombat | 34 | 74 | 4.1 | 3 | 8.9 | 428 | 64 |
| 20 | *Monodelphis domestica* | Gray short-tailed opossum | 34 | 107 | 3.7 | 4 | 9.9 | 468 | 65 |
| 21 | *Varanus komodoensis* | Komodo dragon | 34.5 | 124 | 3.2 | 4 | 11.8 | 626 | 66 |
| 22 | *Elephantulus edwardii* | Cape Rock Elephant Shrew | 35 | 226 | 5.8 | 13 | 10.9 | 580 | 67 |
| 23 | *Sarcophilus harrisii* | Tasmanian devil | 35.5 | 141 | 5.0 | 7 | 11.1 | 491 | 68 |
| 24 | *Fukomys damarensis* | Damaraland mole-rat | 35.5 | 556 | 26.4 | 147 | 6.8 | 953 | 69 |
| 25 | *Phascolarctos cinereus* | Koala | 36 | 113 | 6.2 | 7 | 10.1 | 467 | 70 |
| 26 | *Lipotes vexillifer* | Yangtze river dolphin (baiji) | 36 | 351 | 19.1 | 67 | 7.5 | 705 | 71 |
| 27 | *Loxodonta africana* | African elephant | 36 | 449 | 21.2 | 95 | 7.0 | 804 | 72 |
| 28 | *Notamacropus eugenii* | Tammar wallaby | 36 | 113 | 7.1 | 8 | 10.2 | 467 | 73 |
| 29 | *Tursiops truncatus* | Bottlenose dolphin | 36.5 | 192 | 7.3 | 14 | 8.0 | 547 | 74 |
| 30 | *Homo sapiens* | Human | 36.5 | 428 | 19.4 | 83 | 7.6 | 780 | 75 |
| 31 | *Mus musculus* | Domestic mouse | 36.5 | 332 | 17.2 | 57 | 8.2 | 686 | 17,76 |
| 32 | *Macaca mulatta* | Rhesus macaque | 37 | 413 | 18.9 | 78 | 7.4 | 798 | 77 |
| 33 | *Mirounga angustirostris* | Northern elephant seal | 37 | 374 | 15.0 | 56 | 8.6 | 727 | 78 |
| 34 | *Physeter macrocephalus* | Sperm whale | 37 | 265 | 15.5 | 41 | 9.4 | 619 | 79 |
| 35 | *Myotis lucifugus* | Little brown bat | 37 | 322 | 17.4 | 56 | 9.7 | 678 | 80 |
| 36 | *Equus caballus* | Horse | 37.5 | 466 | 25.6 | 119 | 7.0 | 820 | 81 |
| 37 | *Ursus arctos* | Brown bear | 37.5 | 343 | 5.5 | 19 | 9.7 | 696 | 82 |
| 38 | *Lontra canadensis* | North American river otter | 38 | 486 | 27.2 | 132 | 7.5 | 839 | 83 |
| 39 | *Octodon degus* | Degu | 38.5 | 508 | 13.8 | 70 | 7.1 | 912 | 84 |
| 40 | *Bos taurus* | Domestic cow | 38.5 | 425 | 17.2 | 73 | 7.2 | 759 | 85 |
| 41 | *Canis lupus familiaris* | Domestic dog | 38.5 | 462 | 27.5 | 127 | 6.95 | 897 | 81 |
| 42 | *Capra hircus* | Domestic goat | 39 | 346 | 18.5 | 64 | 7.2 | 701 | 81 |
| 43 | *Capricornis sumatraensis* | Sumatran serow | 38.5 | 112 | 6.25 | 7 | 6.7 | 466 | 86 |
| 44 | *Oryctolagus cuniculus* | European rabbit | 39 | 524 | 21.8 | 114 | 7.0 | 1022 | 81 |
| 45 | *Sus scrofa* | Domestic pig | 39 | 366 | 16.9 | 62 | 9.1 | 724 | 81 |
| 46 | *Cavia porcellus* | Guinea pig | 39.5 | 338 | 16.0 | 54 | 9.0 | 688 | 87 |
| 47 | *Ochotona princeps* | American pika | 40 | 309 | 24.0 | 74 | 6.3 | 663 | 88 |

Because fertilization temperature spans a wide evolutionary range and NTD length varies substantially across taxa, we used Bayesian regression to quantify uncertainty and assess the robustness of the temperature–NTD length association (Fig. S2). This analysis supported a strong positive posterior association between fertilization temperature and CatSper1 NTD length, with the posterior distribution of the slope parameter overwhelmingly shifted toward positive values (P(β > 0 | data) > 0.99). Cold-fertilizing species, such as marine invertebrates and fishes (6–23 °C) exhibited short N termini (≈ 9–120 aa), whereas warm-fertilizing species (35–40 °C) displayed substantially expanded NTDs (≈ 200–550 aa), with intermediate temperatures showing intermediate NTD lengths. This relationship was consistent across phylogenetic distances (Fig. 1d, Fig. S2, and Table 1).

CatSper1 NTD’s extraordinary variability suggests it may serve as an evolutionary tuning module, potentially enabling species-specific modulation of CatSper activation in response to reproductive thermal environments. The presence and functional importance of the CatSper1 NTD, that features rich histidine content, has puzzled researchers since its original discovery ^1,9^. While it was initially proposed to play a key role in CatSper pH gating, no experimental evidence has yet confirmed this function. The NTD of CatSper1 represents an extreme example of histidine enrichment, exhibiting the highest histidine density reported among pH-sensitive ion channels (Table 2). In mouse CatSper1, histidine accounts for 9.2% of the full-length protein, whereas the N-terminal domain alone contains approximately 17.2% histidine. This level of enrichment places the CatSper1 N-terminus within the range of classical histidine-rich proteins, such as histidine-rich calcium-binding protein (HRC) and histidine-rich glycoprotein (HRG), which contain approximately 11–12.7% histidine (Table 2).

**Table 2.**
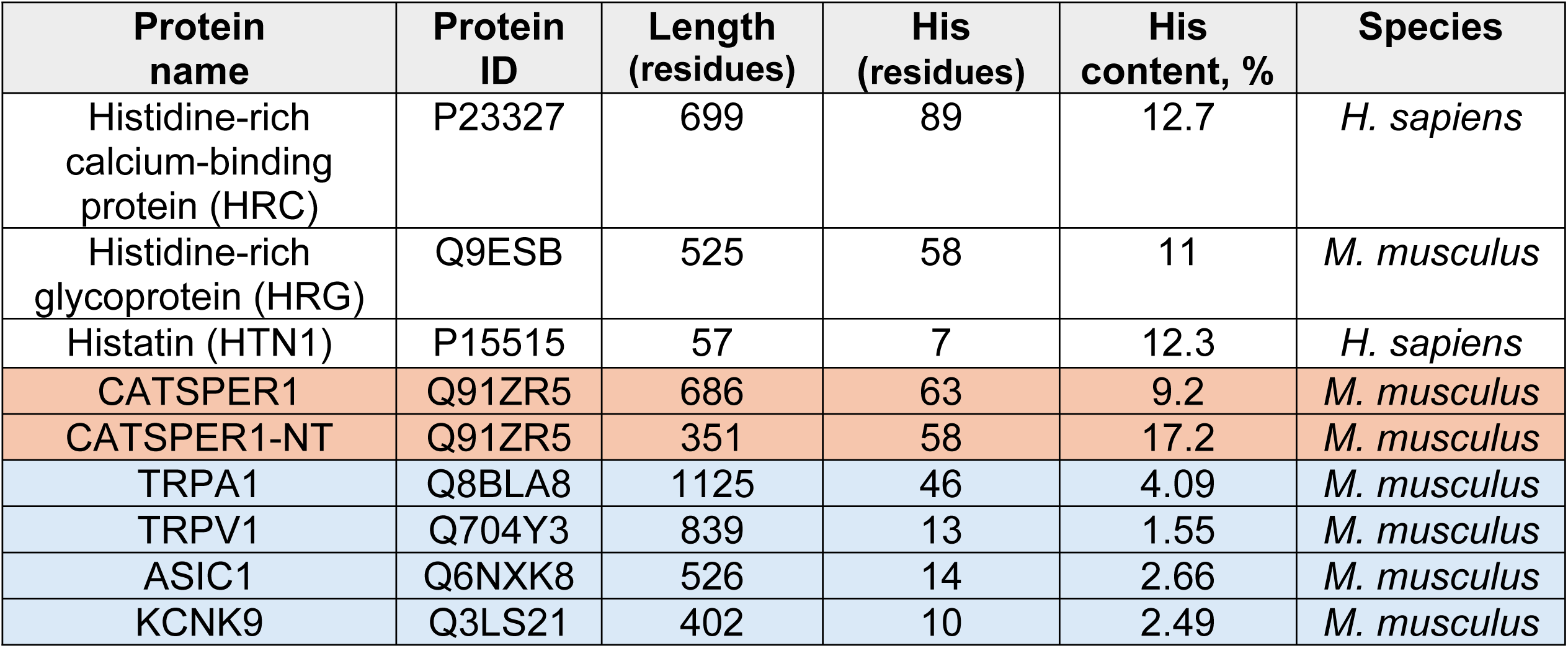
Histidine (His, H) abundance in known histidine-rich proteins. . Histidine-rich proteins are defined by an unusually high abundance of histidine residues. In the average proteome, histidine constitutes approximately 2–3% of amino acids; by contrast, histidine-rich proteins typically contain ≥5–10% histidine, with some domains exhibiting substantially higher levels. Among all pH-sensitive ion channels (blue), only CatSper1 and CatSper N-terminus (CatSper1-NT; red) shows high His content, on par with other known His-rich proteins.

This histidine enrichment of N-terminal region was absent from other pore forming subunits of CatSpermasome, such as CatSper2–4 across the entire evolutionary tree (Fig. 1b and S3a).

In many histidine-rich proteins, clusters of His residues confer sensitivity to pH, metal ions, and other environmental cues, prompting the hypothesis that CatSper1’s N-terminal His clusters contribute to temperature-dependent gating. To test this, we first quantified the total histidine content and found a strong positive correlation with species-specific fertilization temperatures (Fig. 2a), whereas the histidine content of CatSper2–4 showed no correlation (Fig. S3a). Interestingly, an inverse relationship was observed between fertilization temperature and the predicted isoelectric point of the CatSper1 N-terminus (Fig. S3b), consistent with increasing histidine enrichment conferring enhanced pH sensitivity at higher temperatures.

**Figure 2.**
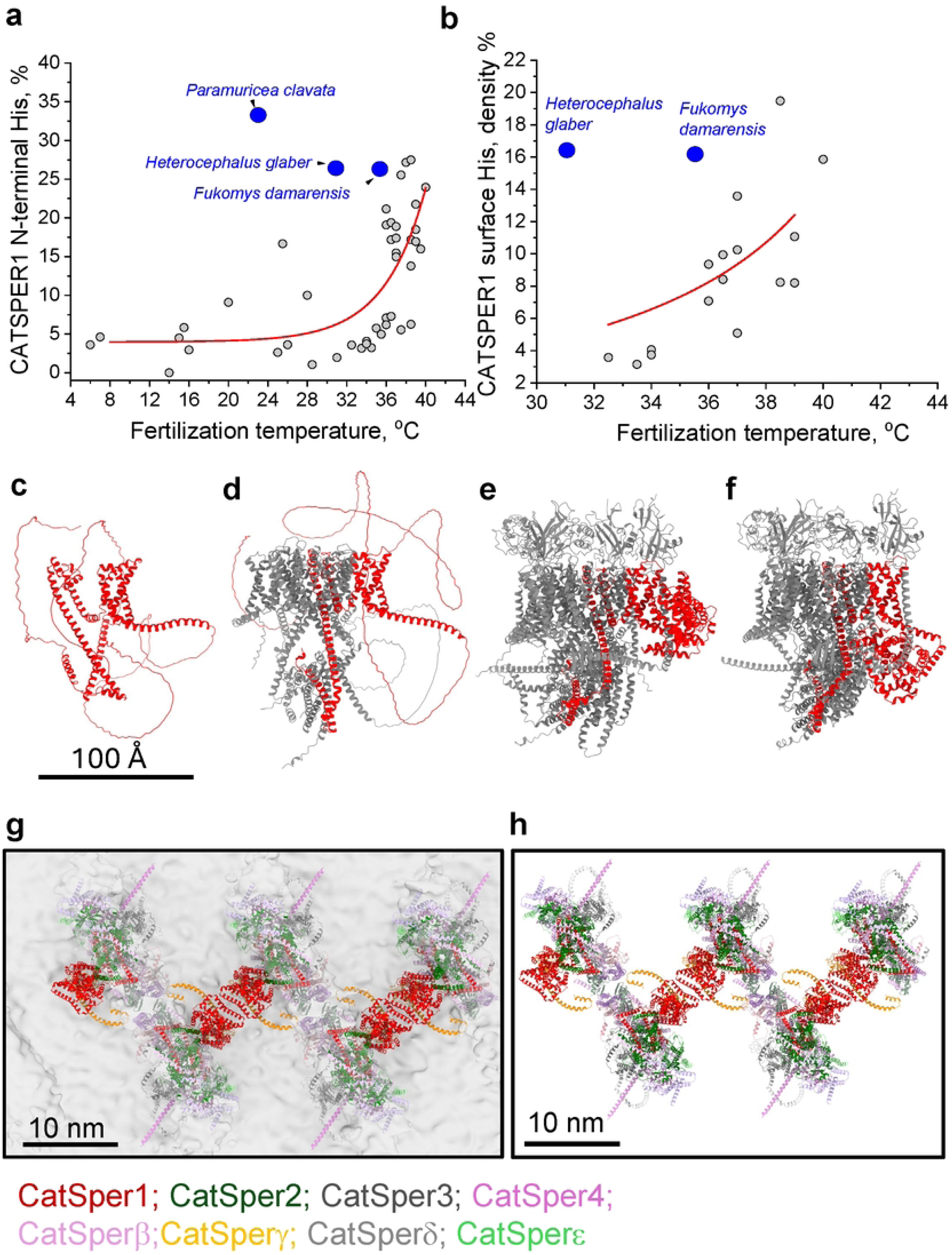
CatSper1 histidine content positively correlates with fertilization temperature. (a) The percentage of histidine residues within the CatSper1 N-terminal domain shows a strong positive correlation with fertilization temperature across CatSper-bearing species shown in Figure 1a. Outliers (blue) correspond to mole rat species, which inhabit subterranean, hypercapnic environments that result in chronically acidic systemic conditions due to elevated carbon dioxide levels. The unusually high apparent percentage of N-terminal histidines in corals *(Paramuricea clavata)* reflects the exceptionally short length of their CatSper1 N-terminal domain (9 residues). (b) Surface His density correlates with fertilization temperatures. (c-f) AlphaFold3 modeling of the murine CatSper1 N-terminal region. (c) The AlphaFold3 predicted structure of a lone murine CatSper1, shown in red. NTD is not predicted to fold into any secondary structures. (d) AlphaFold3 predicted structure of the core subunits CatSper1-4, with CatSper1 shown in red. The NTD is still unstructured. (e) The AlphaFold3 predicted structure of the transmembrane and cytoplasmic parts of the CatSper complex when including intracellular subunits and the transmembrane helices of the extracellular subunits. The NTD is predicted to sit in a region occupied by the cell membrane. (f) The AlphaFold3 predicted structure of the transmembrane and cytoplasmic parts of the CatSper complex as in (e), but with 50 oleic acid molecules added to the prediction. The free oleic acid clusters around the transmembrane domain, causing the NTD to adopt an intracellular position. Each structure is shown with a 100 Å scalebar. (g) Integrative structural model of multiple adjacent murine CatSpermasomes fitted into the cryo-ET density of murine CatSpermasome from ^21^. (h) Same as (g) in the absence of Cryo-ET density. Single-particle cryo-EM structures and AlphaFold3-predicted models were combined to visualize the arrangement of pore-forming and auxiliary subunits within the array. Subunits are color-coded as indicated: CatSper1 (red), CatSper2 (green), CatSper3 (gray), CatSper4 (purple), CatSperβ (light pink), CatSperγ (yellow), CatSperδ (dark gray), and CatSperε (light green).

Secondary structure analysis of the CatSper1 NTD indicates that it possesses hallmark features of an intrinsically disordered region (IDR)-a protein segment that lacks a stable tertiary structure and exhibits high conformational flexibility. Consistent with this prediction, recent high-resolution cryo-EM and in situ 3D cryo-ET structures of the murine CatSper complex ^21,31^, obtained at 25 °C, resolved the architecture of most CatSpermasome subunits but failed to visualize the CatSper1 NTD. The absence of defined density for this region further supports the notion that the NTD is intrinsically flexible and may participate in dynamic inter-CatSpermasome interactions.

In addition, the CatSper1 NTD displays a high predicted droplet-promoting propensity (pDP score approaching 1.0), suggesting a strong likelihood of liquid–liquid phase separation behavior. High pDP values (∼1.0) are characteristic of proteins capable of droplet formation and self-assembly into higher-order condensates, processes that are often sensitive to environmental factors such as temperature and ionic strength. Thus, phase separation mediated by the CatSper1 IDR may contribute to the distinctive regulatory properties of CatSper1 (Fig. S4).

To determine whether the CatSper1 IDR can adopt a stable three-dimensional structure under physiological conditions, we integrated AlphaFold3 predictions with available structural data. Specifically, we modeled the murine CatSper1 NTD in isolation, in the context of neighboring CatSpermasome components, and within a lipid membrane environment to assess whether molecular context promotes structural stabilization (Fig. 2c-f).

### Comparative genomics combined with AlphaFold3 modeling further supported a relationship between CatSper1 surface-exposed histidine residues and fertilization temperature

As shown in Fig. 2c, initial modeling of CatSper1 NTD in isolation did not predict the formation of defined secondary structural elements, consistent with its intrinsically disordered nature. However, when the transmembrane and cytoplasmic components of the CatSper complex were included, comprising the intracellular subunits and the transmembrane helices of extracellular subunits, the NTD was positioned within a region corresponding to the plasma membrane interface (Fig. 2d-e). Upon further incorporation of free oleic acid molecules clustering around the transmembrane domain to better approximate a lipid environment, the NTD adopted an intracellular orientation (Fig. 2f). This positioning is consistent with its proposed localization between adjacent CatSpermasomes^32,38,39^ and enabled quantitative assessment of the density of surface-exposed histidine residues within the NTD. The calculated histidine surface density was then plotted against species-specific fertilization temperatures, revealing a strong positive correlation across mammals (Fig. 2b).

Two exceptions—*Heterocephalus glaber* and *Fukomys damarensis*—possess extremely high N-terminal His content (∼26%) yet intermediate core body temperatures (30–36 °C; Fig. 2a-b, Fig. S5 and Table 1). These subterranean mole rats inhabit hypoxic, CO₂-rich burrows, likely imposing unique selective pressures on sperm physiology. Such an environment may have contributed to lineage-specific adaptations in CatSper channel function and regulation, setting these species apart from other mammals in the evolutionary trajectory of sperm motility control.

To identify a potential interaction module between CatSper1 N-termini, we fitted the high-resolution cryo-EM structure of the CatSpermasome ^31^ into the density maps of CatSpermasome arrays previously resolved by cryo-electron tomography (cryo-ET^21^).

The predicted intracellular and transmembrane architecture of the CatSper complex was then integrated into the resulting 3D reconstruction (Fig. 2g-h and Fig. 3).

**Figure 3.**
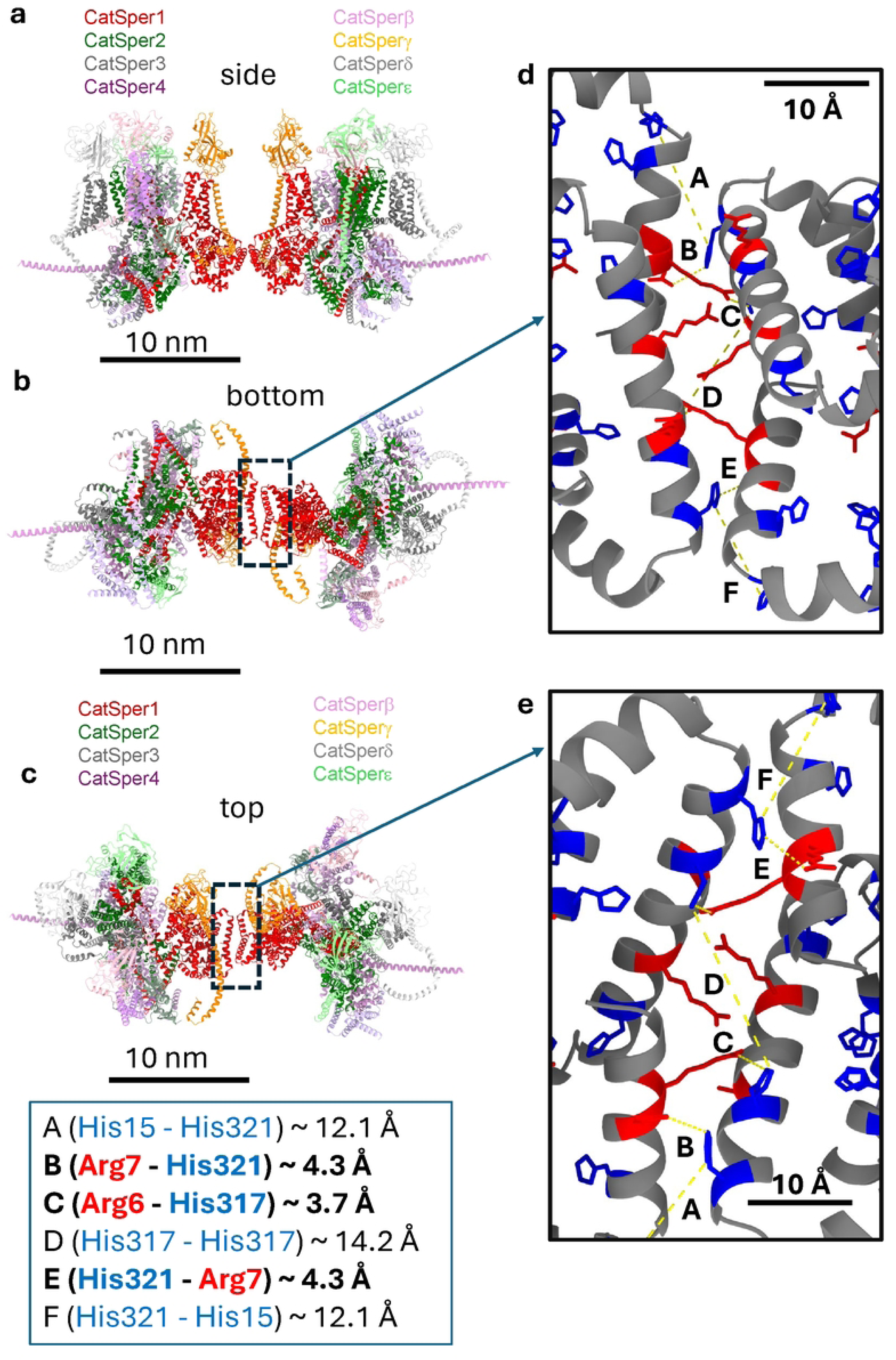
Structural modeling identifies a putative CatSper1 N-terminal interaction interface between neighboring CatSpermasomes. (a–c) Integrative model of two adjacent CatSpermasomes viewed from the **side** (a), **bottom** (b), and **top** (c). Models were generated by fitting the high-resolution single-particle cryo-EM structure of the CatSper complex from ^31^ into the in situ cryo-electron tomography (cryo-ET) density of CatSper arrays from ^21^, followed by incorporation of AlphaFold3-predicted transmembrane and intracellular regions. Pore-forming subunits are color-coded: CatSper1 (red), CatSper2 (green), CatSper3 (gray), and CatSper4 (purple). Dashed boxes indicate the region of closest approach between adjacent CatSpermasomes. (d) Magnified view of the boxed region in (b), highlighting paired α-helical elements from the CatSper1 NTDs positioned at the inter-complex interface. Histidine side chains are shown in blue, arginine side chains are shown in red. Dashed yellow lines indicate distances between conserved His-His residues and His-Arg residues across neighboring complexes: **A**, His15–His321 (∼12.1 Å); **B**, Arg7–His321 (∼4.3 Å); **C**, Arg6–His317 (∼3.7 Å); **D**, His317–His317 (∼14.2 Å); **E**, His321–Arg7 (∼4.3 Å); and **F**, His321–His15 (∼12.1 Å). (e) Magnified view of the boxed region in (c), illustrating the spatial distribution of surface-exposed histidine residues along the interacting helices. Scale bars, 10 nm (a–c) and 1 nm (d-e).

This composite model revealed a previously unrecognized putative interaction interface formed by the proximal regions of CatSper1 N-terminal domains, spanning residues 6-321 in murine CatSper1. These segments are positioned at the junction between adjacent CatSpermasomes within the longitudinal arrays, suggesting a role in inter-complex coupling. AlphaFold3 predicts that this interaction module is composed of paired α-helical elements contributed by opposing CatSper1 N-termini. Each helix contains three conserved histidine residues (H15, H317, and H321; Fig. 3 and Fig. S5). The spatial arrangement of these residues defines a potential interaction network, including contacts between H15 from one CatSpermasome and H321 from an adjucent complex, as well as reciprocal interactions between opposing H317 residues (Fig. 3a-c and S5).

Emerging evidence indicates that histidine residues within intrinsically disordered domains can mediate context-dependent stabilization ^40–42^. In addition to π–π or His–His interactions, neutral histidines at pH values above ∼6.8 can participate in strong cation–π interactions with positively charged residues such as arginine and lysine, promoting local structural ordering. Notably, two arginine residues (Arg6 and Arg7) are located within the predicted murine CatSper1 NTD interface. The modeled distances between these arginines and neighboring histidines (3.7–4.3 Å) fall within the range compatible with stabilizing cation–π interactions. Such proximity suggests a potential mechanism for pH-dependent inter-CatSpermasome coupling mediated by Arg–His interactions (Fig. 3a-c and S5-S6a).

Our modeling is based on the available cryo-ET^21^ map (Fig. 2g) obtained at room temperature (25 °C), which is below the activation threshold of murine CatSper^17^. This likely explains the relatively long distances observed between putative interacting histidines (H317–H317 and H15–H321), which range from ∼11.8–14.2 Å—too large to support direct π–π or hydrogen-bond interactions. We therefore propose that elevated temperature promotes histidine deprotonation, reducing electrostatic repulsion and enabling closer association of CatSper1 NTDs. Consistent with this idea, phylogenetic analysis reveals that H317 (in murine CatSper1) is the most conserved histidine residue within this interface, with notable divergence in cold-blooded internal fertilizers such as reptiles, supporting a role in temperature-dependent regulation (Fig. 3 and S5).

Together, these findings identify a histidine-arginine-based interface that may mediate pH-and temperature-dependent coupling between CatSpermasomes.

Histidine-rich proteins are known to serve diverse regulatory functions, including pH buffering, metal ion coordination, and modulation of protein stability. Importantly, the pKa of histidine (∼6.3)^43^ allows it to readily switch between charged and neutral states within a narrow physiological pH range (6.5–7.5), which coincides with the conditions that regulate CatSper activity.

In temperature-sensitive proteins, increased temperature can shift local pKa values, weakening hydrogen bonding networks and promoting histidine deprotonation. Conversely, under acidic conditions, histidine protonation leads to charge accumulation that can destabilize protein structure by introducing electrostatic repulsion and disrupting hydrophobic packing. In fact, as we have shown recently, cytoplasmic acidification to 6.0 profoundly attenuated CatSper’s temperature gating^17^, which further supports CatSper N-terminal involvement in this process.

### Experimental evidence indicates that removal of CatSper1 N-terminus results in the loss of temperature-dependent gating

In our recent work, we demonstrated that murine spermatozoa subjected to prolonged capacitation (>60 minutes) progressively lose temperature-dependent gating of CatSper^17^. This loss is accompanied by a time-dependent decline in CatSper-mediated currents for both divalent (Ba²⁺) and monovalent (Cs⁺) ions (Fig. 4a and ^17^). Using direct sperm patch-clamp recordings, we further observed a progressive reduction in temperature-dependent sodium conductance through CatSper during prolonged sperm capacitation (Fig. 4b).

**Figure 4.**
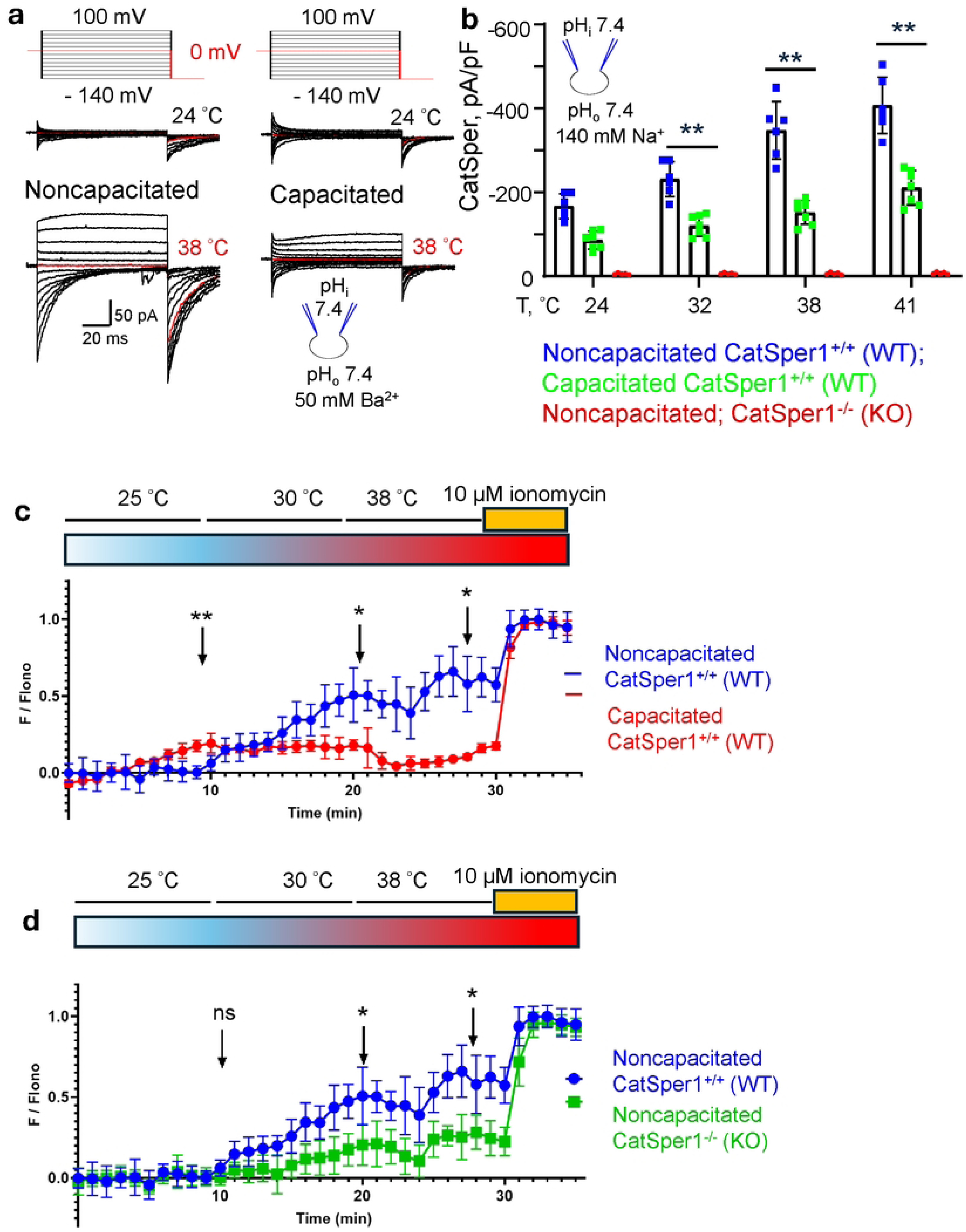
Sperm capacitation leads to removal of temperature sensitivity. (a) Representative divalent Ba^2+^ currents (*I_Ba_^2+^*) recorded from wild type noncapacitated mouse spermatozoa (left) and from wild type capacitated sperm cell (right) in response to indicated voltage steps under two different temperatures: 24 °C and 38 °C. Exposure to 38 °C increases *I_Ba_^2+^* stronger in noncapacitated cells. (b) Decrease in current density of Na^+^ currents via CatSper with sperm capacitation after response to elevated temperatures. CatSper1^-/-^ sperm were used as negative control. Data are mean values ± S.E.M., n corresponds to the number of cells used; n=5-7. (c) CatSper plays a critical role in temperature-regulated increases in sperm [Ca^2+^]_i_. Average traces of normalized Fluo4-AM fluorescence in wild-type noncapacitated mouse spermatozoa (blue) and from capacitated sperm (red) in response to increasing temperatures: 25°C, 30°C, and 38°C. (d) Average traces of normalized Fluo4-AM fluorescence in noncapacitated wild-type (blue) and CatSper1^-/-^ (green) mouse spermatozoa in response to increasing temperatures: 25°C, 30°C, and 38 °C. Each trace was normalized to its respective ionomycin (Ionomycin, 10 μM) response. Data are presented as mean and standard deviation (n=3 biological replicates). *P<0.05, **P<0.01, ns=non-significant, by unpaired t-test.

To independently validate these findings, we performed calcium imaging experiments to assess CatSper activity in response to its threshold temperature activation in non-capacitated versus capacitated sperm. In agreement with electrophysiological recordings, capacitated sperm failed to exhibit temperature-dependent Ca²⁺ influx. CatSper1-deficient sperm were used as controls to exclude non–CatSper-mediated effects. As shown in Fig. 4c-d, capacitation completely abolished temperature-dependent activation of murine CatSper.

Prior studies from Chung et al and Ded et al. ^14,36^ convincingly demonstrated that these capacitating conditions induce selective proteolytic removal of the CatSper1 N-terminus^14,36^, temporally coinciding with the loss of temperature gating observed in our experiments (Fig. 4). Together, these findings support the conclusion that temperature gating is critical for maintaining CatSper channels organized within CatSpermasomes in a highly ordered and synchronized array, enabling their concerted opening. Disruption of this domain uncouples channels from the ordered CatSpermasome lattice, leading to asynchronous or inefficient channel opening. Loss of concerted gating reduces effective channel conductance and limits Ca²⁺ influx, thereby impairing the robust calcium signaling required to drive hyperactivated sperm motility.

Based on these principles, we propose that the high histidine content of the CatSper1 N-terminal domain paired with adjacent positively charged residues (Arg, Lys) confers sensitivity to temperature-dependent histidine deprotonation, stabilizing inter-CatSpermasome interactions in a manner analogous to alkaline pH. In contrast, intracellular acidification would promote histidine protonation, destabilizing the N-terminal interface and inhibiting channel activity (Fig. 5). This model is fully consistent with experimental observations showing that CatSper is activated by warm temperatures (>34 °C)^17^ and alkaline pH^6,44^, while acidic pH potently suppresses channel function.

**Figure 5.**
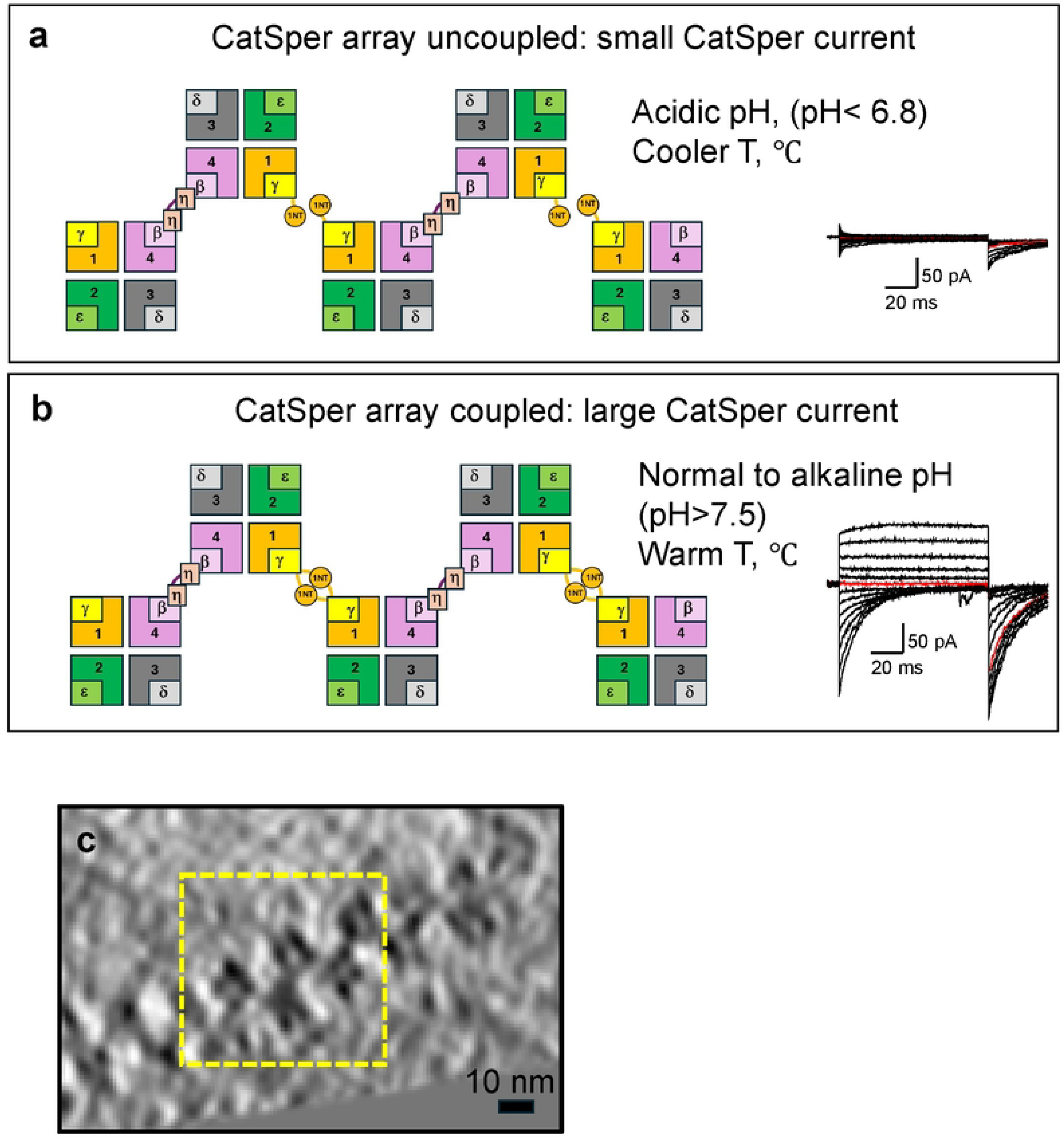
Model for pH-and temperature-dependent coupling of CatSpermasomes mediated by the CatSper1 N-terminus. (a) Schematic representation of CatSpermasome organization within the zigzag array under acidic intracellular pH and/or cooler temperature conditions. Under these conditions, protonation of histidine residues within the CatSper1 NTDs (1NT) is proposed to weaken 1NT interactions and reduce cooperative channel coupling: array uncoupled. Right insert: representative small CatSper current is shown. (b) Corresponding schematic demonstrates array coupling under alkaline intracellular pH and warm temperature conditions. Deprotonation of histidine residues will ensure histidine-arginine coupling, stabilizing interactions between neighboring CatSpermasomes, and promoting cooperative coupling along the array. Representative enlarged CatSper current is shown on the right. (c) Representative cryo-electron tomography (cryo-ET) slice of the human CatSper channel array in the sperm flagellum from ^21^, viewed parallel to the principal piece axis. The dashed yellow box highlights a repeating CatSpermasome unit within the longitudinal zigzag array.

According to this model, pH-and temperature-dependent deprotonation of surface-exposed histidine residues within the CatSper1 N-termini, including His317 and His321, stabilizes cation–π interactions with Arg6 and Arg7, and promotes coupling between adjacent CatSpermasomes, thereby enabling synchronized channel activation at elevated temperatures (Fig. 5).

## DISCUSSION

In addition to regulation by pH and temperature, CatSper activity is strongly modulated by membrane voltage, such that progressively greater depolarization is required to open the channel under acidic conditions ^6,17,19,44,45^. Conversely, at warmer temperatures, CatSper opens at less depolarized potentials, whereas acidic pH completely abolishes temperature-dependent gating. This interdependence between temperature, pH, and voltage suggests a shared structural basis for polymodal regulation.

We propose that this integration is mediated by the spatial proximity of the CatSper1 N-terminal interaction module to the S4 segment of the CatSper1 voltage-sensor domain (VSD). The S4 segment of CatSper1 is the strongest voltage-sensing element among the pore-forming subunits, harboring an unusually high density of positively charged residues—four arginines (Arg; R) and two lysines (Lys; K)—that together form a classical voltage-gated motif (RxxRxxRxxR; Fig. S6b and ^9,31^). This charge density is predicted to confer heightened voltage sensitivity relative to CatSper2–4, whose S4 segments contain progressively fewer charged residues, with CatSper4 harboring only two Arg and one His. These structural differences point to a hierarchical organization of gating contributions, with CatSper1 acting as the dominant voltage sensor and the remaining subunits serving modulatory roles.

The close apposition of the CatSper1 VSD and NTD suggests that these elements function as a coupled gating unit, integrating voltage sensing with temperature-and pH-dependent regulation. Upon membrane depolarization, upward movement of the S4 helix is expected to mechanically displace the adjacent NTD. Based on the predicted organization of CatSpermasome arrays, neighboring CatSper channels may interact through their NTD, potentially via coiled-coil–like associations. In this configuration, depolarization-induced S4 movement would impose mechanical tension on the N-terminal interface, promoting coordinated conformational transitions and priming adjacent channels for synchronized opening.

Inter-CatSpermasome coupling interfaces have previously been identified within the canopy region and at the interface between the CatSper4 and CatSperβ subunits ^38,39^. However, the interface formed by the CatSper1 N-terminal domain (NTD) has remained unresolved and was largely considered an intrinsically disordered region with unknown functional properties. To our knowledge, this is the first report assigning a specific structural and functional role to the CatSper1 NTD in coordinating channel opening within longitudinal CatSper arrays.

Recent quantum chemical analyses by Calinsky et al ^40–42^ have demonstrated that histidine residues within intrinsically disordered domains can mediate pH-dependent structural assembly through dual π–π and cation–π interactions. At physiologically relevant alkaline pH, deprotonated (neutral) histidine can form stabilizing cation–π interactions with positively charged residues such as arginine (Arg) and lysine (Lys). Notably, two arginine residues are present within the predicted interface between CatSper1 N-termini, supporting the possibility of such stabilizing interactions under alkaline conditions.

In contrast, at acidic pH, protonated histidine preferentially engages in cation–π interactions with aromatic residues (e.g., Tyr, Phe, Trp), interactions that are typically stronger than those formed by neutral histidine with cationic residues. Importantly, aromatic residues are absent from the predicted CatSper1 NTD interface, suggesting that acidification would not promote alternative stabilizing contacts.

Although the arginine residues predicted to mediate cation–π interactions within the murine CatSper1 NTD appear to be poorly conserved across warm-fertilizing species, their functional role may be substituted by lysine residues (Fig. S6a). Lysines are more broadly conserved among these species and are positioned in close proximity to the predicted interface region, making them plausible alternative partners for stabilizing cation–π interactions with histidines. Nevertheless, high-resolution atomic structures of CatSper complexes from additional species will be required to directly test and validate this hypothesis.

Together, these findings identify a histidine-dependent coupling interface positioned adjacent to the dominant voltage-sensing element of the CatSpermasome, providing a structural framework for the interplay between temperature, pH, and voltage in CatSper gating. In this model, acidic pH is predicted to disrupt N-terminal coupling by protonating histidine residues within the interface, increasing electrostatic repulsion and weakening inter-channel interactions. This would reduce coupling efficiency, leading to asynchronous channel opening and diminished cumulative Ca²⁺ influx. Conversely, elevated temperature and alkaline pH would favor histidine deprotonation, stabilize N-terminal interactions, and enhance cooperative gating within CatSper arrays (Fig. 5).

More broadly, this work establishes CatSper as a new model for how evolutionary pressures sculpt ion-channel architecture to integrate environmental and physiological signals essential for optimal sperm performance.

This model is grounded in comparative genomics, structural modeling, and integration of high-resolution cryo-EM and cryo-ET data with functional analysis of murine sperm cells. However, definitive validation of the proposed histidine-dependent coupling mechanism will require atomic resolution of CatSper conformational dynamics across defined pH, temperature, and voltage conditions. In particular, high-resolution structural snapshots captured under physiologically relevant temperatures and targeted mutagenesis of conserved histidine residues will be essential to resolve the precise molecular rearrangements underlying synchronized gating. Such experiments are beyond the scope of the present study but are a natural and necessary extension of this work, and we anticipate that future experimental structures will further refine and strengthen the mechanistic model proposed here.

## METHODS

### Comparative genomics & sequence alignment

Protein sequences for CatSper subunits and related proteins across a wide range of species were retrieved using Python Scripts accessing the UniProtKB, UniParc, and NCBI APIs. Data extraction was done in June 2025. Sequences labeled as partial or containing the names of other subunits were removed. In cases where a single species had multiple isoforms, the longest sequence was selected. Suspected misannotated sequences were visually identified in an alignment and replaced using NCBI Protein BLAST searches with the corresponding mouse subunit. Species with at least 4 pore-forming subunits (Catsper1-4) conserving pore-forming subunits (Catsper1-4) were chosen for sequence alignment. An evolutionary tree was built using NCBI Common Tree [PMID: 32761142].

### Definition of the CatSper1 N-terminal region

To define the boundaries of the Catsper1 N-terminal region, all full-length Catsper1 protein sequences from the dataset were aligned using MUSCLE (v5.2)^89^ with default parameters. The resulting alignment was used to identify the start of the conserved pore-forming region, defined as the first position in the alignment containing a block of at least 10 consecutive columns with greater than 80% occupancy. For each individual sequence, the non-gap characters in the alignment preceding this position were used as the N-terminal region. The N-terminal sequence spanned from the beginning of the sequence to the beginning of the short 20 amino acid (murine) linker region connecting the N-terminal domain to the S1 transmembrane domain.

### Quantification of N-Terminal histidine content

To calculate N-terminal histidine abundance, the number of histidine residues in an N-terminal sequence was normalized to the total number of amino acids in the N-terminus.

### Computation of physicochemical properties of the CatSper1 N-terminus

Theoretical isoelectric points for each N-terminal sequence were calculated using the Expasy ProtParam web server ^90^. Liquid-Liquid Phase Separation (LLPS) probabilities were calculated using the FuzDrop ^91^ web server, using the mouse N-terminal sequence as a representative model. A pDP score greater than 0.6 was used as the threshold for identifying likely LLPS regions.

### Catspermasome structural modeling and bioinformatic analysis

Structural models of the transmembrane and intracellular parts of the CatSper complex were generated using AlphaFold3 ^35^ In cases where model inference failed, auxiliary subunits distant from the N-terminal region were excluded to improve model inference. The last 280 residues of each of the extracellular subunit sequences, containing the transmembrane helices, were included in the AlphaFold3 prediction as separate chains. Fifty oleic acid ligands were added to the Alphafold3 predictionsprediction to mimic the membrane environment. The N-terminal domain was only predicted to fold into secondary structures when the transmembrane helices of the extracellular subunits were included in the prediction (Figure S5). Five models were generated for each complex. For each species, the model with the highest-ranking score as calculated by Alphafold3 was used for all further analysis.

Single-particle cryo-EM maps (EMD-31076) ^31^ were fitted as rigid bodies into the cryo-ET structure for the in situ arrangement of the CatSper complex (EMD-24210)^21^ using ChimeraX ^92^. Alphafold3 predicted structures for the transmembrane and intracellular regions of the CatSper complex were fitted to these cryo-EM maps to visualize the relative orientations of complexes. A residue was defined as being on the surface if its Relative Solvent Accessibility (RSA) was greater than 25%. RSA was calculated for each residue using the formula RSA = (SASA / MaxSASA) * 100. Only residues in the N-terminal region were accounted for in the calculation. The Solvent Accessible Surface Area (SASA) was calculated for each PDB model using the Shrake-Rupley algorithm as implemented in the Bio.PDB.SASA package in Biopython (v1.85)^93^. Maximum SASA values (MaxSASA) for each amino acid were taken from the reference values published by Tien et al. (2013)^94^.

### Bayesian regression analysis

To assess the relationship between fertilization temperature and CatSper1 N-terminal (NT) length across species, we performed a Bayesian regression analysis using data summarized in Table 1 and Fig. 1d. NT length was modeled as a shifted exponential function (NT_i_) of fertilization temperature:

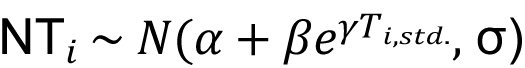

where 𝑇𝑖,𝑠𝑡𝑑. represents the standardized temperature 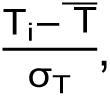 represents the baseline NT length, 𝛽 represents the scaling parameter of exponential growth, 𝛾 represents the growth rate parameter, and 𝜎 represents the residual variance. Priors were chosen to be weakly informative and biologically plausible, reflecting observed NT length ranges:

𝛼 ∼ 𝑁(100,50),

𝛽 ∼ 𝐻𝑎𝑙𝑓𝑁𝑜𝑟𝑚𝑎𝑙(50),

𝛾∼𝐿𝑜𝑔𝑁𝑜𝑟𝑚𝑎𝑙(0,1),

𝜎 ∼ HalfNormal(50).

Posterior distributions were estimated by Markov chain Monte Carlo sampling using the No-U-Turn Sampler implemented in PyMC^95^. Four chains were run for 3000 iterations, each following 2000 tuning steps with a target acceptance rate of 0.95. Inference was based focused on the posterior probability distributions of all parameters.

### Animals and ethics statement

All mouse procedures were performed according to the National Institutes of Health Guiding Principles for the care and use of laboratory animals. These procedures were reviewed and approved by the Institutional Animal Care and Use Committee of Washington University in St. Louis (St. Louis, MO, USA) (protocol # 20-0126 and protocol 22-0251). C57BL/6, C57BL/6NJ and CatSper1 knock-out male mice (CatSper KO) (60–90 days old) were purchased from Charles River and Jackson Laboratory (JAX) and kept at a constant temperature of 22 ± 2 °C under a 12/12 h dark/light cycle with free access to food and water at the barrier animal facility of WashU School of Medicine. The animals were fed a standard chow diet. There was no difference in the sperm quality between mouse lines. CatSper1^-/-^ males were born at a Mendelian ratio by breeding CatSper1^-/-^ females with CatSper1^+/-^ males. Genotypes were continuously monitored by TransnetYX, Inc. (TN) and verified by in-house genotyping as reported in^17^.

### Mouse sperm collection and capacitation

Sexually mature male mice (60–90 days old) were euthanized using CO₂ followed by cervical dislocation, and sperm were isolated from the cauda epididymis. Sperm were allowed to swim-up in non-capacitating (NC) TYH media buffered with HEPES (in mM: 119.3 NaCl, 4.7 KCl, 1.71 CaCl_2_.2H_2_O, 1.2 KH_2_PO_4_, 1.2 MgSO_4_.7H_2_O, 25.1 NaHCO_3_, 5.56 glucose, 0.51 sodium pyruvate, 10 HEPES) at pH 7.4 for 15–20 min at 37 °C, unless specified. The motile fraction of the sample was removed from the tube. To induce capacitation, sperm were incubated in capacitating (CAP) TYH medium supplemented with 5 mg/mL BSA and 15 mM NaHCO_3_ at 37 °C for 60-90 min. NC sperm were incubated in TYH without BSA and NaHCO_3_.

### Sperm intracellular calcium concentration measurement

After sperm swim-up in NC or CAP conditions for 20 min, motile sperm were collected and incubated with 10 μM Fluo-4 AM and 0.05% Pluronic F-127 at 37 °C for 60-90 min. Sperm were centrifuged at 300-400 g for 10 min and further resuspended in the corresponding media. Cells (5X10^6^/well) were seeded in a 384-well black, clear-bottom plate (Greiner Bio-One) and sperm [Ca^2+^]_i_ was recorded at 25, 30, and 38 °C using a CLARIOstar plate reader (BMG LABTECH). The instrument was set to bottom-reading mode with an excitation wavelength of 483-14 nm and emission of 530-30 nm. Acquisition parameters used were: 10 flashes per well with a cycle time of 30 sec per read. Gain was adjusted to 50% on the highest signal well. The protocol was set to double orbital shaking at 300 rpm for 10 s before each measurement. Data was collected with MARS Data Analysis software. For each condition, an initial 10-minute baseline reading at 25 °C was followed by sequential 10-minute recordings at 30 and 38 °C, followed by a final perfusion with 10 μM Ionomycin (Iono) at the end of every experiment. Clampfit 10 (Molecular Devices), and GraphPad were used for data analysis. To compare fluorescence across different samples and experimental conditions, Fluo-4 fluorescence (F) in response to different temperatures for each sperm was normalized to its respective F_Iono_ (F/F_iono_).

## Acknowledgements

This work was supported by BJC Investigator fund to P.V.L., startup funding from the McDonnell Center for Cellular and Molecular Neurobiology, the Department of Cell Biology & Physiology, and the Center for the Investigation of Membrane Excitability Diseases at Washington University in St. Louis to ZF.

---

**Figure S1.**
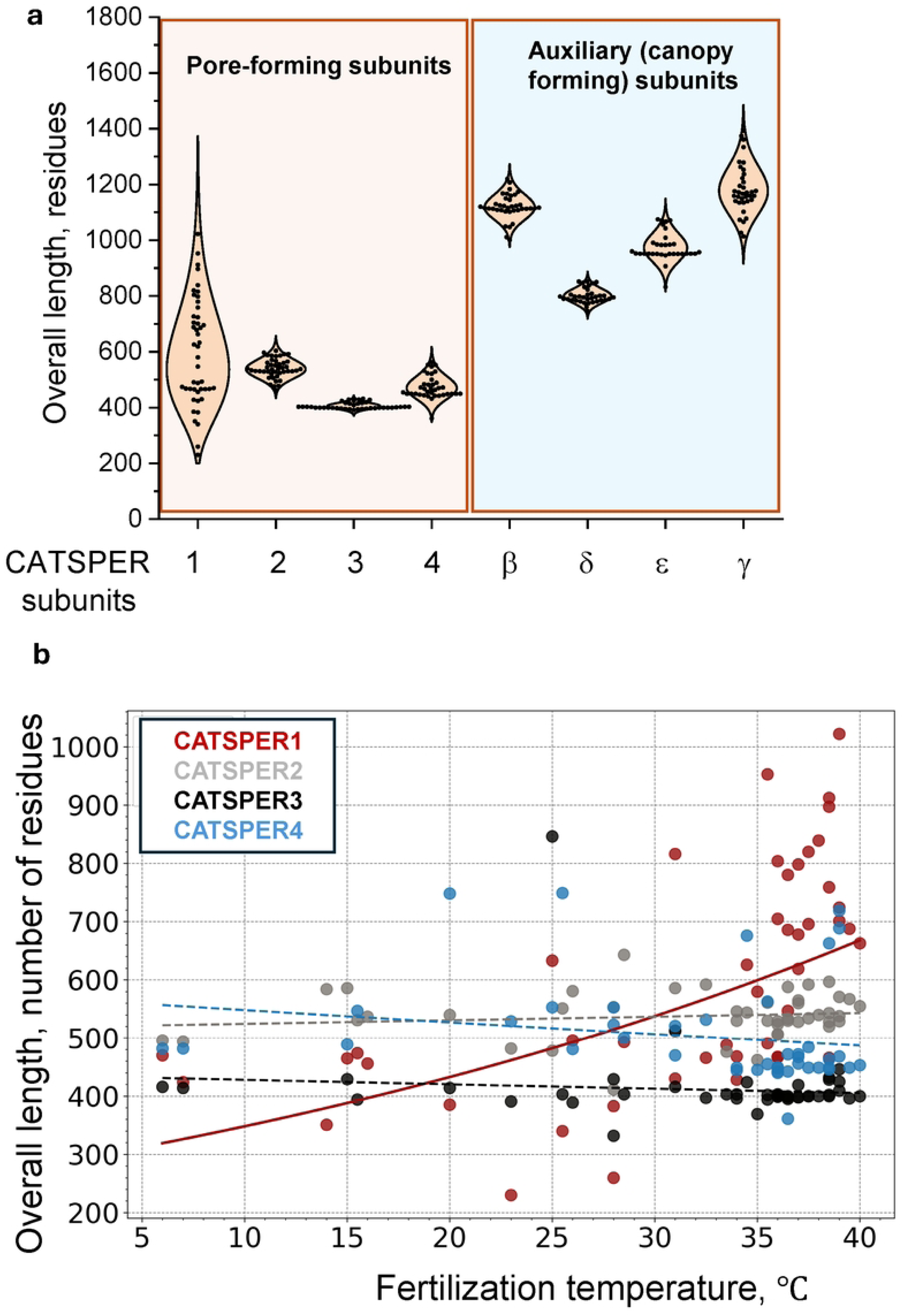
Comparative genomic analyses of CatSpermasome transmembrane subunits. (a) Comparison of the total amino acid length of eight CatSpermasome core-forming subunits across species shown on Figure 1a. (b) CatSper1-4 entire length plotted against fertilization temperature for the species shown in Figure 1a.

**Figure S2.**
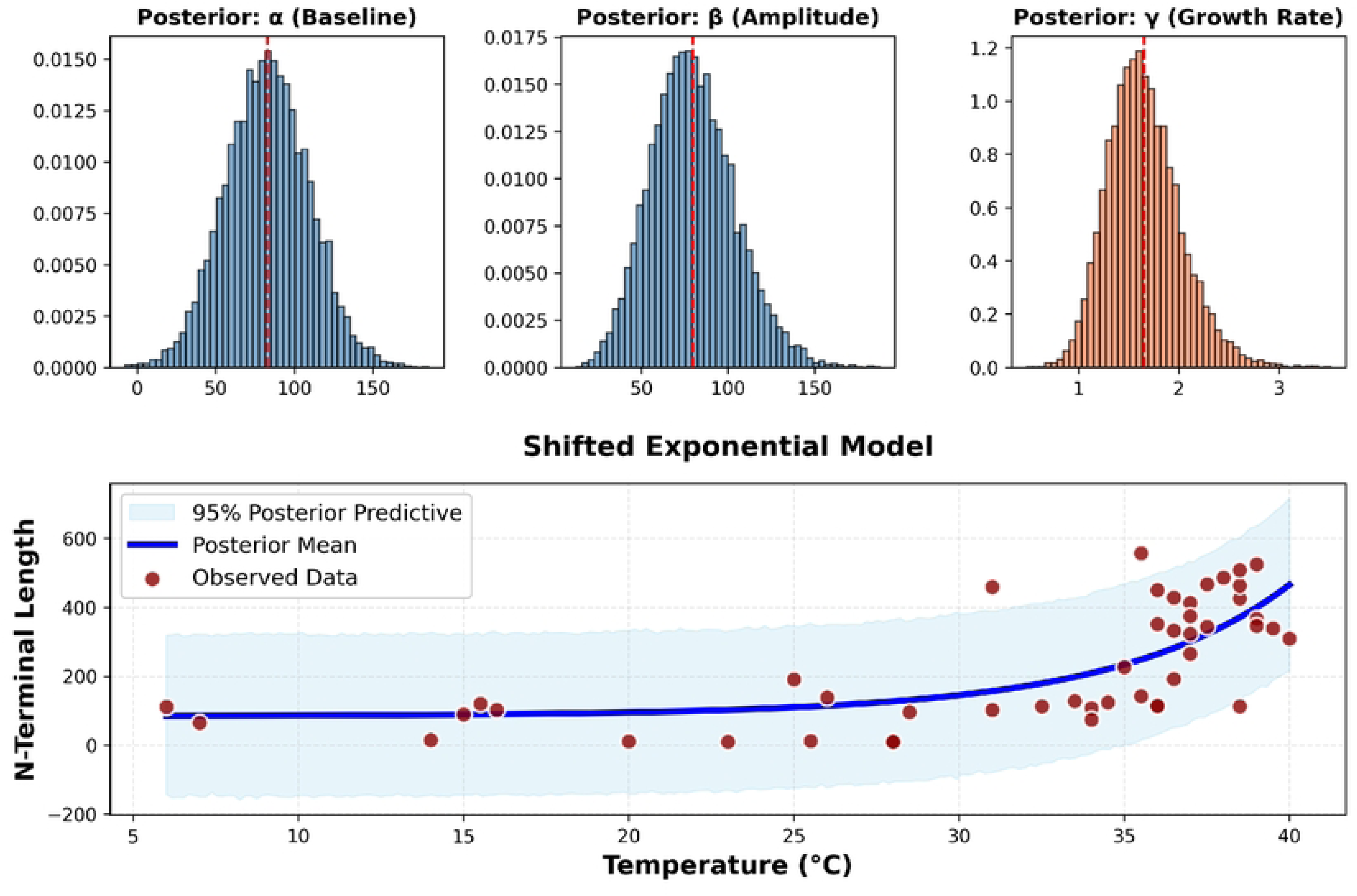
Bayesian regression analysis supports strong positive posterior association between fertilization temperature and CatSper1 N-terminal length. Priors were chosen to be weakly informative and biologically plausible, reflecting observed NT length ranges: 𝛼 ∼ 𝑁(100,50); 𝛽 ∼ 𝐻𝑎𝑙𝑓𝑁𝑜𝑟𝑚𝑎𝑙(50); 𝛾∼𝐿𝑜𝑔𝑁𝑜𝑟𝑚𝑎𝑙(0,1), 𝜎 ∼ 𝐻𝑎𝑙𝑓𝑁𝑜𝑟𝑚𝑎𝑙(50). Parameters were inferred using Monte Carlo sampling. Four chains were run for 3000 iterations, each following 2000 tuning steps with a target acceptance rate of 0.95. Trace and posterior plots show convergence of the chains and the distribution of model parameters.

**Figure S3.**
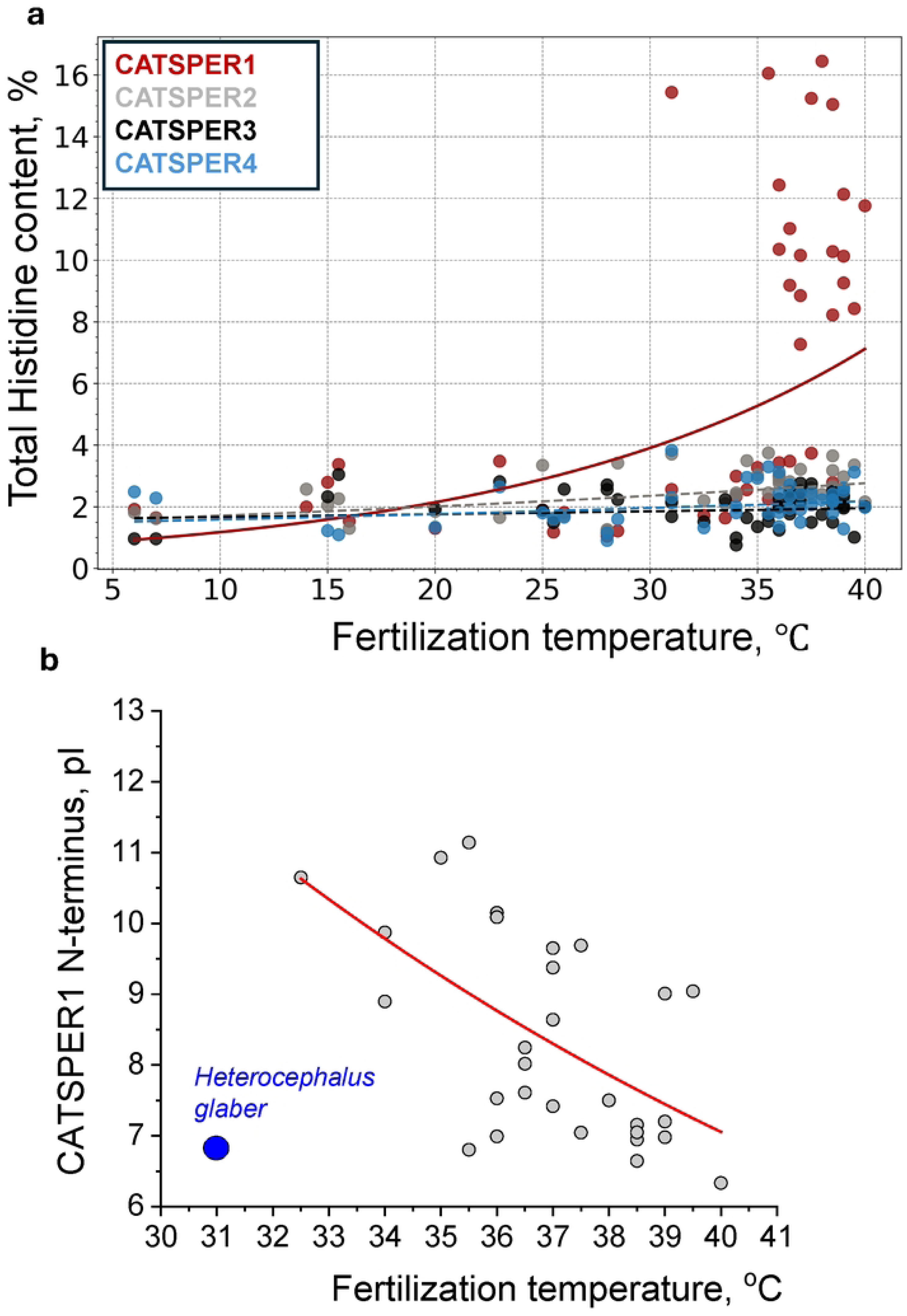
Correlation between CatSper1 N-terminal histidine content, pI and fertilization temperatures. (a) The percentage of histidine residues within entire CatSper1-4 subunits plotted against fertilization temperature. Only CatSper1 shows positive correlation. (b) Theoretical isoelectric point (pI) calculated for N-terminus domain of CatSper1 plotted against fertilization temperature for mammalian species shown in Figure 1a.

**Figure S4.**
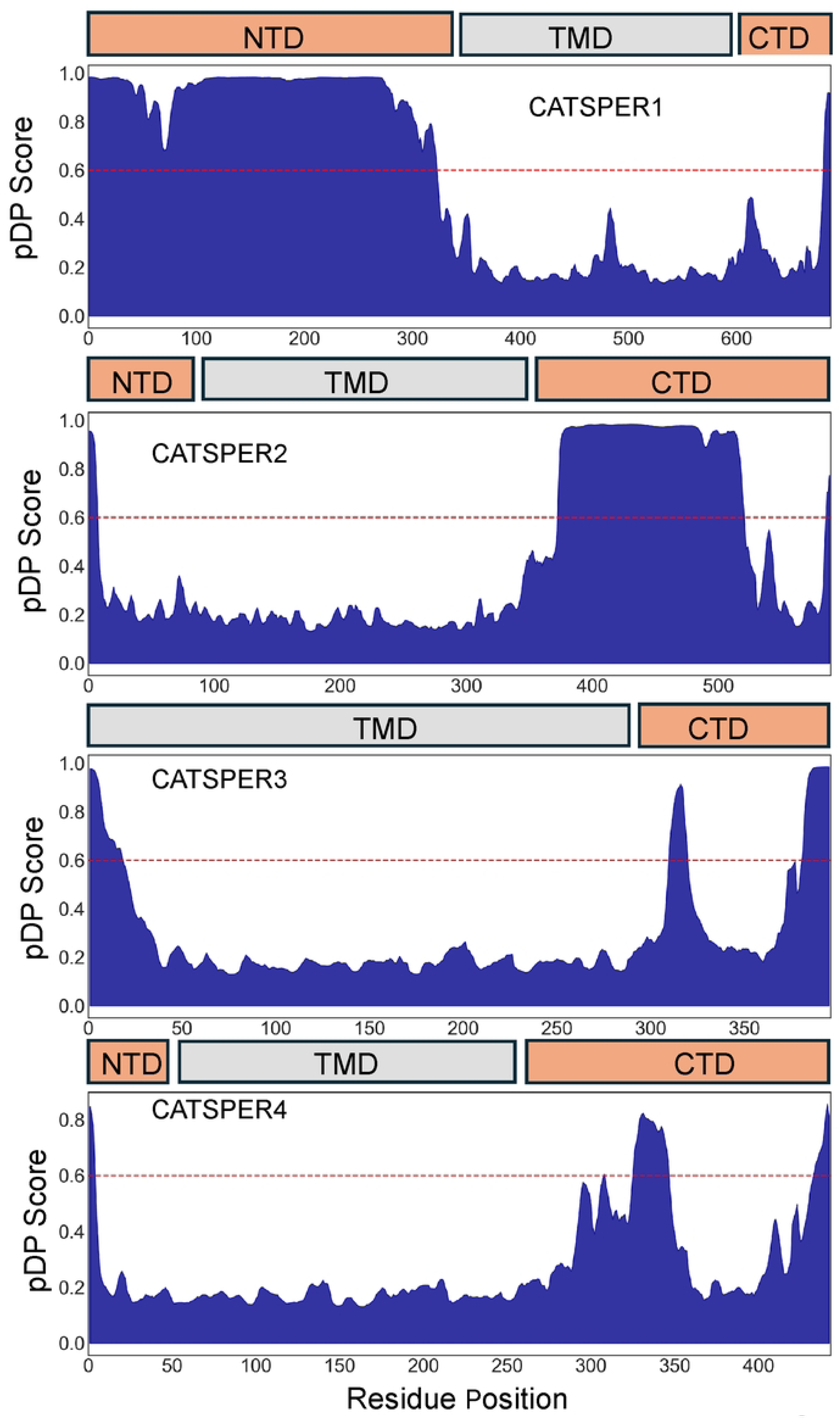
The CatSper1 N-terminal domain uniquely exhibits high predicted liquid–liquid phase separation propensity. Predicted liquid–liquid phase separation (LLPS) propensity of CatSper pore-forming subunits. pDP scores calculated using FuzDrop are plotted as a function of residue position for CatSper1, CatSper2, CatSper3, and CatSper4. Domain boundaries are indicated schematically above each plot: N-terminal domain (NTD, peach), transmembrane domain (TMD, gray), and C-terminal domain (CTD, peach). The red dashed line denotes the pDP threshold of 0.6, above which regions are predicted to have a high propensity for droplet formation.CatSper1 displays an extended N-terminal region with consistently high pDP scores exceeding the LLPS threshold, whereas CatSper2–4 lack comparable LLPS-prone regions in their N-terminal domains and instead show elevated pDP scores primarily within C-terminal regions. These results indicate that a predicted intrinsically droplet-forming region is a unique feature of the CatSper1 N-terminus among the pore-forming CatSper subunits.

**Figure S5.**
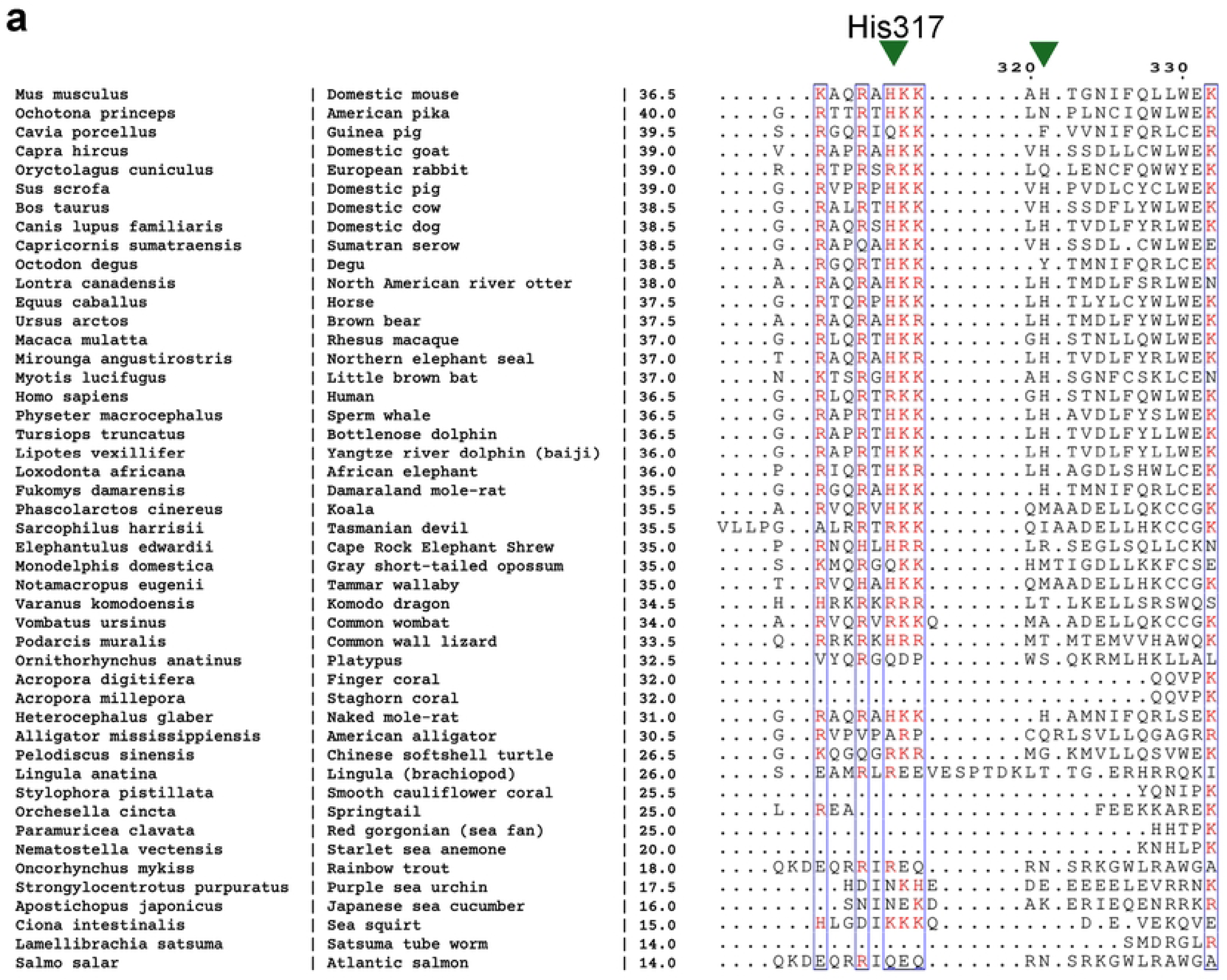
Sequence alignment of CatSper1 NTDs between residues 300 and 330 of murine CatSper1 across species. Conserved His residues indicated by arrowheads.

**Figure S6.**
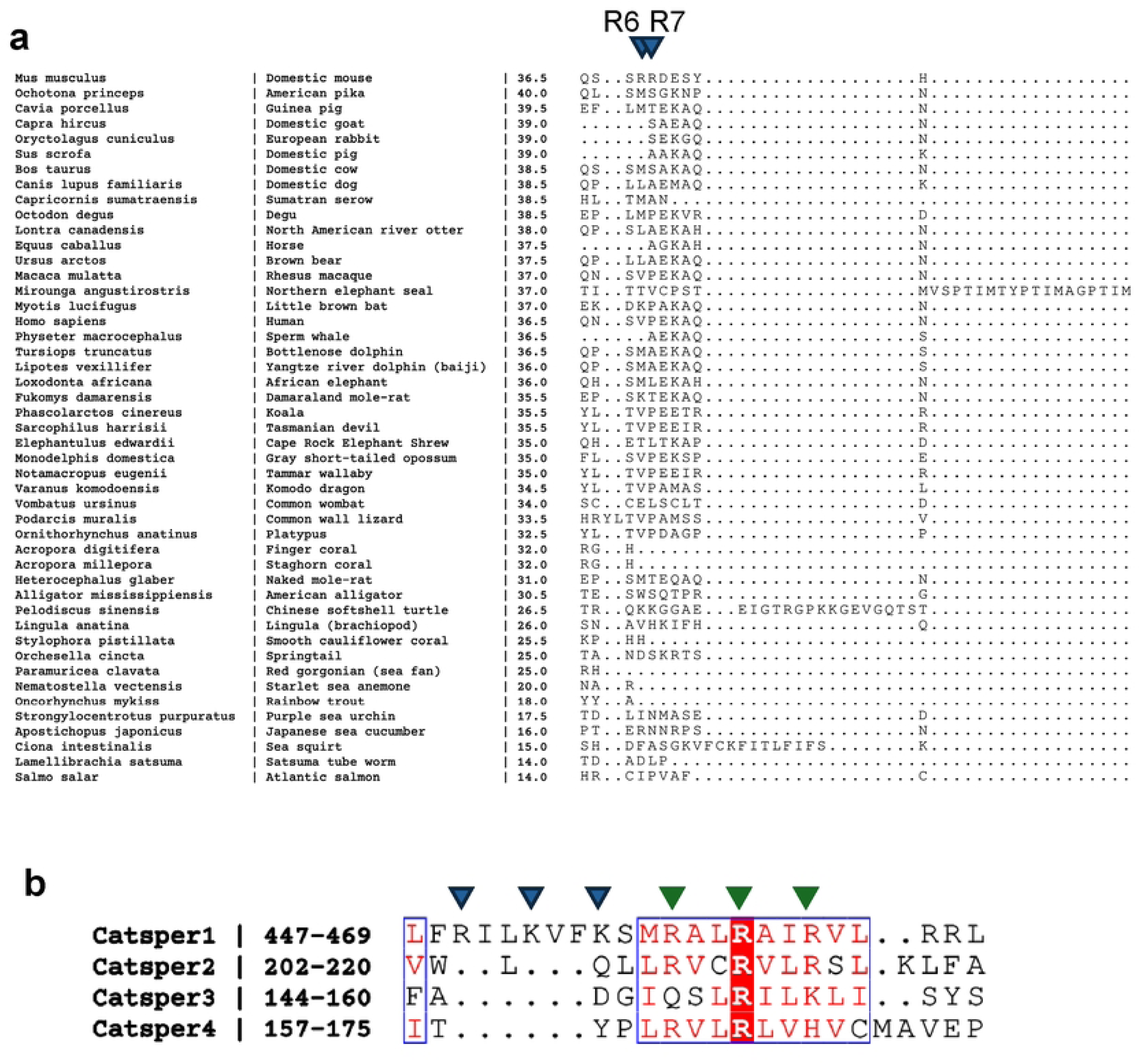
Hierarchical organization of CatSper voltage-gated module located in S4 regions. (a) Sequence alignment of CatSper1 NTDs withing first 10 resides across 47 species. (b) Sequence alignment between S4 regions of murine CatSper1 to 4. An unusually high density of positively charged residues, four R and two K, is located in the S4 region of CatSper1, which is predicted to enable enhanced voltage sensitivity. CatSper2–4, however, contain progressively fewer charged residues, with CatSper4 harboring only two R. Conserved positively charged residues are indicated by green triangles. Blue triangles indicate additional residues unique to CatSper1.

## References

1. Ren, D., et al. A sperm ion channel required for sperm motility and male fertility. Nature 413, 603–609 (2001).

2. Quill, T.A., Ren, D., Clapham, D.E. & Garbers, D.L. A voltage-gated ion channel expressed specifically in spermatozoa. Proc Natl Acad Sci U S A 98, 12527–12531 (2001).

3. Carlson, A.E., et al. CatSper1 required for evoked Ca2+ entry and control of flagellar function in sperm. Proc Natl Acad Sci U S A 100, 14864–14868 (2003).

4. Lobley, A., Pierron, V., Reynolds, L., Allen, L. & Michalovich, D. Identification of human and mouse CatSper3 and CatSper4 genes: characterisation of a common interaction domain and evidence for expression in testis. Reprod Biol Endocrinol 1, 53 (2003).

5. Avidan, N., et al. CATSPER2, a human autosomal nonsyndromic male infertility gene. Eur J Hum Genet 11, 497–502 (2003).

6. Kirichok, Y., Navarro, B. & Clapham, D.E. Whole-cell patch-clamp measurements of spermatozoa reveal an alkaline-activated Ca2+ channel. Nature 439, 737–740 (2006).

7. Qi, H., et al. All four CatSper ion channel proteins are required for male fertility and sperm cell hyperactivated motility. Proc Natl Acad Sci U S A 104, 1219–1223 (2007).

8. Suarez, S.S. Control of hyperactivation in sperm. Hum Reprod Update 14, 647–657 (2008).

9. Navarro, B., Kirichok, Y., Chung, J.J. & Clapham, D.E. Ion channels that control fertility in mammalian spermatozoa. Int J Dev Biol 52, 607–613 (2008).

10. Cai, X. & Clapham, D.E. Evolutionary genomics reveals lineage-specific gene loss and rapid evolution of a sperm-specific ion channel complex: CatSpers and CatSperbeta. PLoS One 3, e3569 (2008).

11. Hildebrand, M.S., Avenarius, M.R. & Smith, R.J.H. CATSPER-Related Male Infertility. (2009).

12. Lishko, P.V. & Kirichok, Y. The role of Hv1 and CatSper channels in sperm activation. J Physiol 588, 4667–4672 (2010).

13. Chung, J.J., Navarro, B., Krapivinsky, G., Krapivinsky, L. & Clapham, D.E. A novel gene required for male fertility and functional CATSPER channel formation in spermatozoa. Nat Commun 2, 153 (2011).

14. Chung, J.J., et al. Structurally distinct Ca(2+) signaling domains of sperm flagella orchestrate tyrosine phosphorylation and motility. Cell 157, 808–822 (2014).

15. Cai, X., Wang, X. & Clapham, D.E. Early evolution of the eukaryotic Ca2+ signaling machinery: conservation of the CatSper channel complex. Molecular biology and evolution 31, 2735–2740 (2014).

16. Quill, T.A., et al. Hyperactivated sperm motility driven by CatSper2 is required for fertilization. Proc Natl Acad Sci U S A 100, 14869–14874 (2003).

17. Swain, D.K., Vergara, C., Castro-Arnau, J. & Lishko, P.V. The essential calcium channel of sperm CatSper is temperature-gated. Nat Commun 16, 3657 (2025).

18. Lishko, P.V., Botchkina, I.L., Fedorenko, A. & Kirichok, Y. Acid extrusion from human spermatozoa is mediated by flagellar voltage-gated proton channel. Cell 140, 327–337 (2010).

19. Lishko, P.V., Botchkina, I.L. & Kirichok, Y. Progesterone Activates the Principal Ca2+ Channel of Human Sperm. Nature 471, 387–391 (2011).

20. Hwang, J.Y. & Chung, J.J. CatSper Calcium Channels: 20 Years On. Physiology (Bethesda) 38, 0 (2023).

21. Zhao, Y., et al. 3D structure and in situ arrangements of CatSper channel in the sperm flagellum. Nat Commun 13, 3439 (2022).

22. Hwang, J.Y., et al. Dual Sensing of Physiologic pH and Calcium by EFCAB9 Regulates Sperm Motility. Cell 177, 1480–1494 e1419 (2019).

23. Chung, J.J., et al. CatSperzeta regulates the structural continuity of sperm Ca2+ signaling domains and is required for normal fertility. eLife 6(2017).

24. Brenker, C., et al. The CatSper channel: a polymodal chemosensor in human sperm. EMBO J 31, 1654–1665 (2012).

25. Wang, H., Liu, J., Cho, K.H. & Ren, D. A novel, single, transmembrane protein CATSPERG is associated with CATSPER1 channel protein. Biol Reprod 81, 539–544 (2009).

26. Ho, K., Wolff, C.A. & Suarez, S.S. CatSper-null mutant spermatozoa are unable to ascend beyond the oviductal reservoir. Reproduction, fertility, and development 21, 345–350 (2009).

27. Liu, J., Xia, J., Cho, K.H., Clapham, D.E. & Ren, D. CatSperbeta, a novel transmembrane protein in the CatSper channel complex. J Biol Chem 282, 18945–18952 (2007).

28. Xia, J., Reigada, D., Mitchell, C.H. & Ren, D. CATSPER channel-mediated Ca2+ entry into mouse sperm triggers a tail-to-head propagation. Biol Reprod 77, 551–559 (2007).

29. Jin, J., et al. Catsper3 and Catsper4 are essential for sperm hyperactivated motility and male fertility in the mouse. Biol Reprod 77, 37–44 (2007).

30. Carlson, A.E., et al. Identical phenotypes of CatSper1 and CatSper2 null sperm. J Biol Chem 280, 32238–32244 (2005).

31. Lin, S., Ke, M., Zhang, Y., Yan, Z. & Wu, J. Structure of a mammalian sperm cation channel complex. Nature 595, 746–750 (2021).

32. Zhao, Q., et al. ARMH2 is a cytosolic component of CatSper crucial for sperm function. Nat Commun 16, 10243 (2025).

33. Huang, X., et al. A CUG-initiated CATSPERtheta functions in the CatSper channel assembly and serves as a checkpoint for flagellar trafficking. Proc Natl Acad Sci U S A 120, e2304409120 (2023).

34. Cai, X. & Clapham, D.E. Ancestral Ca2+ signaling machinery in early animal and fungal evolution. Molecular biology and evolution (2011).

35. Abramson, J., et al. Accurate structure prediction of biomolecular interactions with AlphaFold 3. Nature 630, 493–500 (2024).

36. Ded, L., Hwang, J.Y., Miki, K., Shi, H.F. & Chung, J.J. 3D in situ imaging of the female reproductive tract reveals molecular signatures of fertilizing spermatozoa in mice. Elife 9(2020).

37. Brown, S.G., et al. Homozygous in-frame deletion in CATSPERE in a man producing spermatozoa with loss of CatSper function and compromised fertilizing capacity. Hum Reprod 33, 1812–1816 (2018).

38. Hwang, J.Y., et al. CATSPERepsilon extracellular domains are essential for sperm calcium channel assembly and activity modulation. Sci Adv 12, eadw3414 (2026).

39. Xu, Q., et al. Molecular basis of the higher-order assembly of CatSper. Proc Natl Acad Sci U S A 123, e2510754123 (2026).

40. Calinsky, R. & Levy, Y. Aromatic Residues in Proteins: Re-Evaluating the Geometry and Energetics of pi-pi, Cation-pi, and CH-pi Interactions. The journal of physical chemistry. B 128, 8687–8700 (2024).

41. Calinsky, R. & Levy, Y. Histidine in Proteins: pH-Dependent Interplay between pi-pi, Cation-pi, and CH-pi Interactions. Journal of chemical theory and computation 20, 6930–6945 (2024).

42. Calinsky, R. & Levy, Y. A pH-Dependent Coarse-Grained Model for Disordered Proteins: Histidine Interactions Modulate Conformational Ensembles. The journal of physical chemistry letters 15, 9419–9430 (2024).

43. Bashford, D. & Karplus, M. pKa’s of ionizable groups in proteins: atomic detail from a continuum electrostatic model. Biochemistry 29, 10219–10225 (1990).

44. Lishko, P.V., et al. The control of male fertility by spermatozoan ion channels. Annu Rev Physiol 74, 453–475 (2012).

45. Lishko, P., Clapham, D.E., Navarro, B. & Kirichok, Y. Sperm patch-clamp. Methods in enzymology 525, 59–83 (2013).

46. Dodge, D.P. & Maccrimmon, H.R. Environmental Influences on Extended Spawning of Rainbow Trout (Salmo gairdneri). Transactions of the American Fisheries Society 100, 312–318 (1971).

47. Peterson, R.H., Spinney, H.C.E. & Sreedharan, A. Development of Atlantic Salmon (Salmo salar) Eggs and Alevins Under Varied Temperature Regimes. Journal of the Fisheries Research Board of Canada 34, 31–43 (1977).

48. Southward, Andersen, A. & Hourdez, S. Lamellibrachia anaximandri n. Sp., a new vestimentiferan tubeworm (Annelida) from the Mediterranean, with notes on frenulate tubeworms from the same habitat. Zoosystema 33, 245–279 (2011).

49. Clutton, E.A., Alurralde, G. & Repolho, T. Early developmental stages of native populations of Ciona intestinalis under increased temperature are affected by local habitat history. The Journal of experimental biology 224(2021).

50. Hammond, L.M. & Hofmann, G.E. Thermal tolerance of Strongylocentrotus purpuratus early life history stages: mortality, stress-induced gene expression and biogeographic patterns. Mar Biol 157, 2677–2687 (2010).

51. Yuan, X., Yang, H., Wang, L., Zhou, Y. & Gabr, H.R. Bioenergetic responses of sub-adult sea cucumber Apostichopus japonicus (Selenka) (Echinodermata: Holothuroidea) to temperature with special discussion regarding its southernmost distribution limit in China. Journal of Thermal Biology 34, 315–319 (2009).

52. Stefanik, D.J., Friedman, L.E. & Finnerty, J.R. Collecting, rearing, spawning and inducing regeneration of the starlet sea anemone, Nematostella vectensis. Nature protocols 8, 916–923 (2013).

53. Gómez-Gras, D., et al. Exploring the response of a key Mediterranean gorgonian to heat stress across biological and spatial scales. Scientific reports 12, 21064 (2022).

54. Van Dooremalen, C., Koekkoek, J. & Ellers, J. Temperature-induced plasticity in membrane and storage lipid composition: Thermal reaction norms across five different temperatures. Journal of Insect Physiology 57, 285–291 (2011).

55. Putnam, H.M., Edmunds, P.J. & Fan, T.Y. Effect of temperature on the settlement choice and photophysiology of larvae from the reef coral Stylophora pistillata. The Biological bulletin 215, 135–142 (2008).

56. Emig, C.C. ECOLOGY OF INARTICULATED BRACHIOPODS. Treatise on Invertebrate Paleontology; Part H, 473–526 (1997).

57. Manullang, C., et al. Slight thermal stress exerts genetic diversity selection at coral (Acropora digitifera) larval stages. BMC Genomics 26, 36 (2025).

58. dela Cruz, D.W. & Harrison, P.L. Optimising conditions for in vitro fertilization success of Acropora tenuis, A. millepora and Favites colemani corals in northwestern Philippines. Journal of Experimental Marine Biology and Ecology 524, 151286 (2020).

59. Service, U.S.F.W. Chinese Softshell Turtle (Pelodiscus sinensis) Ecological Risk Screening Summary (ed. Web Version) (2018).

60. Ferguson, M.W. & Joanen, T. Temperature of egg incubation determines sex in Alligator mississippiensis. Nature 296, 850–853 (1982).

61. Riccio, A.P. & Goldman, B.D. Circadian rhythms of body temperature and metabolic rate in naked mole-rats. Physiology & behavior 71, 15–22 (2000).

62. Smyth, D.M. Temperature regulation in the platypus, Ornithorhynchus anatinus (Shaw). Comp Biochem Physiol A Comp Physiol 45, 705–715 (1973).

63. Spears, S., et al. Lizards in the wind: The impact of wind on the thermoregulation of the common wall lizard. Journal of Thermal Biology 121, 103855 (2024).

64. Descovich KA, J.S., Lisle A, Nicolson V, Janssen T, Brooks P, Phillips CJC. Long-term measurement of body temperature in the southern hairy-nosed wombat (Lasiorhinus latifrons). Australian Mammalogy 39, 48–55 (2017).

65. Busse, S., Lutter, D., Heldmaier, G., Jastroch, M. & Meyer, C.W. Torpor at high ambient temperature in a neotropical didelphid, the grey short-tailed opossum (Monodelphis domestica). Naturwissenschaften 101, 1003–1006 (2014).

66. Harlow, H.J., Purwandana, D., Jessop, T.S. & Phillips, J.A. Body temperature and thermoregulation of Komodo dragons in the field. Journal of Thermal Biology 35, 338–347 (2010).

67. Boyles, J.G., Smit, B., Sole, C.L. & McKechnie, A.E. Body temperature patterns in two syntopic elephant shrew species during winter. Comparative biochemistry and physiology. Part A, Molecular & integrative physiology 161, 89–94 (2012).

68. Jones, M.E., Grigg, G.C. & Beard, L.A. Body temperatures and activity patterns of Tasmanian devils (Sarcophilus harrisii) and eastern quolls (Dasyurus viverrinus) through a subalpine winter. Physiol Zool 70, 53–60 (1997).

69. Streicher, S., Boyles, J.G., Oosthuizen, M.K. & Bennett, N.C. Body temperature patterns and rhythmicity in free-ranging subterranean Damaraland mole-rats, Fukomys damarensis. PLoS One 6, e26346 (2011).

70. Adam, D., et al. Body temperature of free-ranging koalas (Phascolarctos cinereus) in south-east Queensland. Int J Biometeorol 64, 1305–1318 (2020).

71. Liu Renjun, Z.X. Artificially rearing the Baiji, Lipotes Vexillifer, for twenty-one years. China Academic Journal Electronic Publishing House (2018).

72. Mole, M.A., Rodrigues, D.S., van Aarde, R.J., Mitchell, D. & Fuller, A. Savanna elephants maintain homeothermy under African heat. J Comp Physiol B 188, 889–897 (2018).

73. Snyder, G.K., Baudinette, R.V. & Gannon, B.J. Oxygen transport and acid-base balance during exercise in the tammar wallaby. Respiration physiology 117, 41–51 (1999).

74. Melero, M., Rodriguez-Prieto, V., Rubio-Garcia, A., Garcia-Parraga, D. & Sanchez-Vizcaino, J.M. Thermal reference points as an index for monitoring body temperature in marine mammals. BMC research notes 8, 411 (2015).

75. Baak, N.A., Cantineau, A.E., Farquhar, C. & Brison, D.R. Temperature of embryo culture for assisted reproduction. The Cochrane database of systematic reviews 9, CD012192 (2019).

76. Walters, E.A., Brown, J.L., Krisher, R., Voelkel, S. & Swain, J.E. Impact of a controlled culture temperature gradient on mouse embryo development and morphokinetics. Reproductive biomedicine online 40, 494–499 (2020).

77. Taffe, M.A. A comparison of intraperitoneal and subcutaneous temperature in freely moving rhesus macaques. Physiology & behavior 103, 440–444 (2011).

78. McGinnis, S.M. & Southworth, T.P. Body Temperature Fluctuations in The Northern Elephant Seal. Journal of Mammalogy 48, 484–485 (1967).

79. Silva, R.G. Assessment of body surface temperature in cetaceans: an iterative approach. Braz J Biol 64, 719–724 (2004).

80. Holyoak, G.W. & Stones, R.C. Temperature regulation of the little brown bat, Myotis lucifugus after acclimation at various ambient temperatures. Comparative Biochemistry and Physiology Part A: Physiology 39, 413–420 (1971).

81. D, R. Temperature Regulation and Thermal Environment, in Dukes’ Physiology of Domestic Animals, (Cornell University Press, 2004).

82. Shah, J., et al. The Brown Bear and Hibernating Mammals as a Translational Model for Human Resilience: Insights for Space Medicine, Critical Care, and Austere Environments. Biology 14, 1434 (2025).

83. North American River Otter. Smithsonian’s National Zoo&Conservation Biology Institute (2025).

84. LafeberVet. The resource for exotic animal veterinary professionals. (2010).

85. Liles, H.L., et al. Positive relationship of rectal temperature at fixed timed artificial insemination on pregnancy outcomes in beef cattle. Journal of animal science 100(2022).

86. Mead, C.N.J.a.J.I. Capricornis crispus. MAMMALIAN SPECIES by the American Society of Mammalogists 750, 1–10 (2004).

87. Namiki, M., Fukayama, T., Suzuki, T. & Masaiwa, A. Relationship between Ear Temperature, Behaviour and Stress Hormones in Guinea Pigs (Cavia porcellus) during Different Interactive Activities in Zoos. Animals (Basel) 14(2024).

88. MacArthur, R.A. & Wang, L.C.H. Physiology of thermoregulation in the pika, Ochotona princeps. Canadian Journal of Zoology 51, 11–16 (1973).

89. Edgar, R.C. MUSCLE: a multiple sequence alignment method with reduced time and space complexity. BMC Bioinformatics 5, 113 (2004).

90. Wilkins, M.R., et al. Protein identification and analysis tools in the ExPASy server. Methods in molecular biology 112, 531–552 (1999).

91. Hardenberg, M., Horvath, A., Ambrus, V., Fuxreiter, M. & Vendruscolo, M. Widespread occurrence of the droplet state of proteins in the human proteome. Proceedings of the National Academy of Sciences 117, 33254–33262 (2020).

92. Goddard, T.D., et al. UCSF ChimeraX: Meeting modern challenges in visualization and analysis. Protein science: a publication of the Protein Society 27, 14–25 (2018).

93. Cock, P.J.A., et al. Biopython: freely available Python tools for computational molecular biology and bioinformatics. Bioinformatics 25, 1422–1423 (2009).

94. Tien, M.Z., Meyer, A.G., Sydykova, D.K., Spielman, S.J. & Wilke, C.O. Maximum allowed solvent accessibilites of residues in proteins. PLoS One 8, e80635 (2013).

95. Abril-Pla, O., et al. PyMC: a modern, and comprehensive probabilistic programming framework in Python. PeerJ Comput Sci 9, e1516 (2023).

